# Attenuated adenosine mediated immune-dampening increases natural killer cell activity in early age-related macular degeneration

**DOI:** 10.1101/2025.01.22.634301

**Authors:** Archana Padmanabhan Nair, Sayan Ghosh, Vishnu Suresh Babu, Machiraju Praveen, Ying Xin, Ganesh Ram Sahu, Tanuja Arun Vaidya, Jayasree Debnath, Karthik Raja, Santhosh Gopi Krishna Gadde, M B Thirumalesh, Naren Shetty, Aishwarya Saxena, Rohit Shetty, Stacey Hose, Vrushali Deshpande, Koushik Chakrabarty, James T. Handa, J. Jiang Qian, Swaminathan Sethu, Debasish Sinha, Arkasubhra Ghosh

## Abstract

Non-exudative age-related macular degeneration (AMD) involves retinal pigment epithelium (RPE) dysfunction and has been linked to altered intraocular immunity. Our investigation focuses on immune cell subsets and inflammation-associated factors in the eyes with early and intermediate AMD. We observed elevated levels of activated natural killer (NK) cells and interferon-γ, concurrent with reduced myeloid-derived suppressor cells (MDSCs) and adenosine in AMD eyes. Aqueous humor from AMD patients had diminished ability to dampen NK cell activation, an effect rescued by adenosine supplementation. The *Cryba1* cKO mouse model recapitulated these immune alterations, and single-cell RNA-sequencing identified NK cell-related genes and NK cell-RPE interactions. Co-culture of activated NK cells with RPE cells induced barrier dysfunction and Gasdermin-E driven pyroptosis providing a functional link relevant to AMD. These findings suggest a double-hit model where elevated immune activation and loss of immune dampening mechanisms drive AMD progression. Resetting the intraocular immune balance may be a promising therapeutic strategy for managing early and intermediate AMD.

**Graphical abstract.**
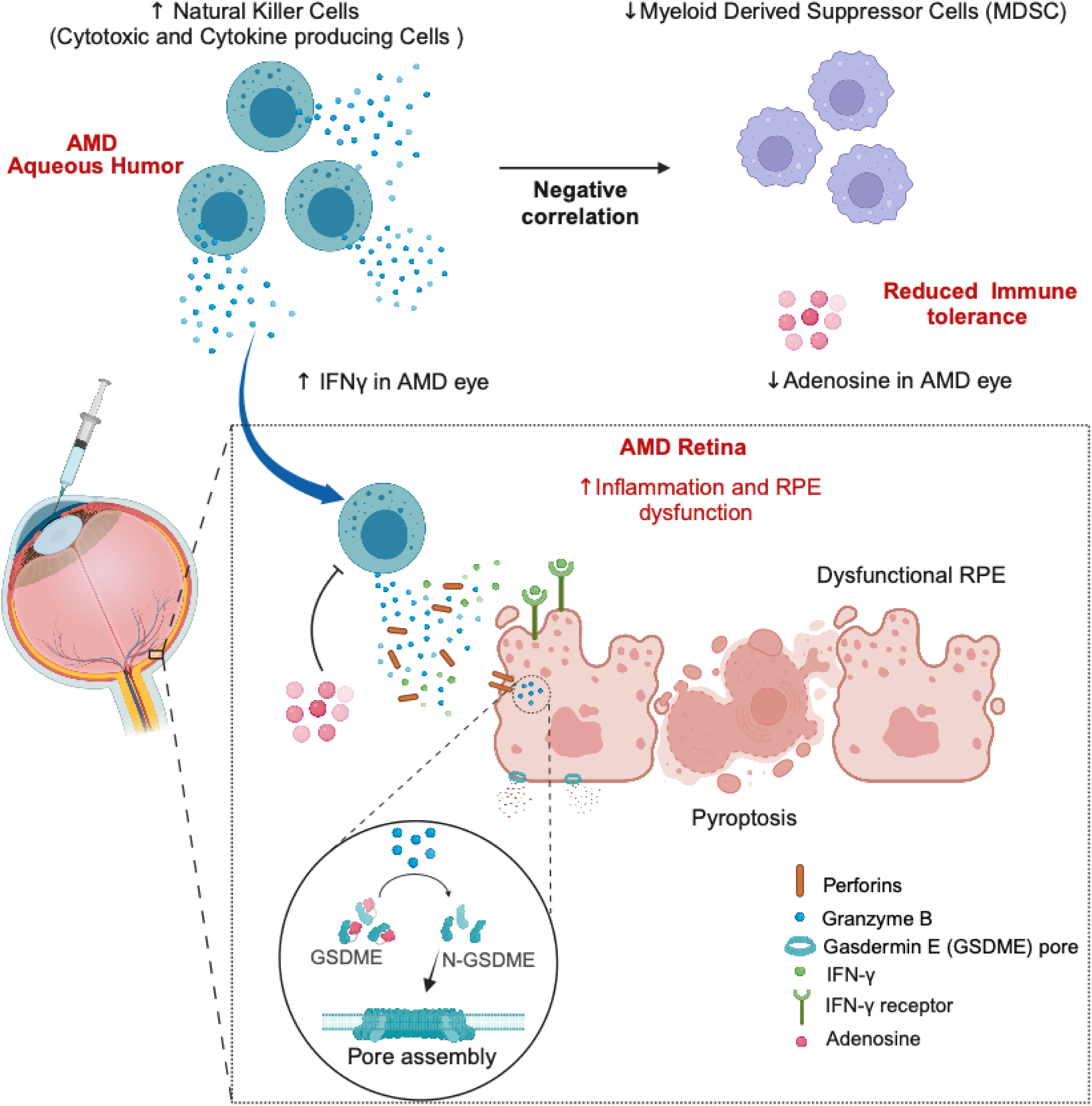
Illustra tion of pro posed mechanism underlying NK cell-RPE interaction in early AMD pathogenesis. Dysregulated NK cell communicate s with stressed RPE in early AMD immunopathology. Aberrant AMD aqueous humor and retina shows increased NK cells and NK effector molecules like IFNγ with reduced MDSC and adenosine in human subjects, don or eye and animal model. Activated NK cells interaction with RPE causes dysfunction and pyroptotic cell death via Gasdermin-E pathway in AMD.. Created with BioRender.com

## INTRODUCTION

Age-related macular degeneration (AMD) is the major contributor of irreversible blindness among the elderly worldwide, currently afflicting 8.8 million people ^1^. Retinal pigment epithelium (RPE) dysfunction and degeneration is a key early change ^2^. Clinically, AMD is diagnosed by loss of central vision, reduced contrast sensitivity and low dark adaptation, metamorphopsia and central scotoma ^3^. The Age-Related Eye Disease Study (AREDS) categorizes AMD severity based on development of Bruch’s membrane deposits called drusen, macular pigmentary alterations resulting in RPE and photoreceptor degeneration, and choroidal thinning into early, intermediate and ultimately, the advanced stages comprised of neovascular AMD (nAMD) or Geographic Atrophy (GA) ^4^. With AMD progression, RPE degeneration can lead to cell death coincident with the development of choriocapillary atrophy and macular photoreceptor death ^5^. The RPE dies by multiple pathways, notably apoptosis in AMD patients, but also inflammasome and amyloid beta fibril mediated pyroptosis ^6,7^, autophagy dysfunction ^8^, and ferroptosis ^9^ have been reported in mouse models. Thus, the mode of RPE cell death may be triggered by the AMD-related stressors it encounters in the milieu.

AMD has multiple risk factors including age, smoking, high fat and glycemic index diets, and genetic predisposition^10^. These factors contribute to key pathogenic pathways including immune dysfunction, oxidative stress, mitochondrial damage, and dysregulated autophagy ^11,12^. Disturbance of both humoral and cellular innate immune systems have been strongly implicated in the AMD pathogenesis ^13–15^. Due to the genetic variants associated with AMD risk, aberrations in the complement system, the humoral arm of the innate immune system, have long been implicated in AMD pathogenesis, and recently, two complement inhibitors have been approved by the FDA for the treatment of geographic atrophy, the advanced dry form of AMD ^16,17^. Besides complement, cells of the innate arm of the immune system such as neutrophils ^18,19^, monocytes ^20–22^, macrophages ^23,24^, mast cells ^25^ and gamma delta T cells ^26^ have also been implicated in AMD. Natural killer (NK) cells, an important member of the innate-lymphoid continuum known to facilitate programmed cell death may be relevant in AMD pathogenesis. Intriguingly, alterations in NK cell phenotypes, subsets, cytokine release, and cytotoxic function have been documented in various age-related conditions, including amyotrophic lateral sclerosis (ALS), Alzheimer’s disease (AD), and stroke ^27^, and perforin/granzyme, a constituent of NK cells, is elevated in AMD retinas and compromises RPE barrier function ^28,29^. However, the role of NK cell mediated effects on RPE dysfunction and pyroptosis in the context of AMD pathogenesis has not been studied.

Ocular immune privilege is an immunosuppressive environment that prevents immune-mediated destruction of ocular cells by the action of immune dampening factors secreted by ocular cells and/or immunoregulatory cells that maintain the blood ocular barrier ^30–35^. In the retina, the outer blood retinal barrier is mediated by the RPE, which is compromised in early AMD ^36^. Notably, secreted anti-inflammatory factors are reduced in the aqueous humor in dry AMD patients, which could disrupt the immunosuppressive microenvironment ^37^. However, status of immunoregulatory cells of the innate arm such as myeloid derived suppressor cells (MDSCs) or factors which have intraocular immune dampening functions are yet to be assessed in AMD patients.

We hypothesized that immune activation and immune dampening mechanisms of the innate arm are imbalanced in AMD, thereby contributing to RPE barrier dysfunction. Studies from various animal models, partly validated in donor human tissues and aqueous humor samples, have implicated a variety of inflammatory factors and immune cell subtypes such as neutrophils, monocytes, macrophages etc. ^38,39^. Yet, detailed studies in early-stage AMD patient eyes are limited, particularly in the context of immune cell repertoire functions. This is primarily due to the paucity of access to human intraocular or retinal samples from early AMD patients and the limited volume of the accessible samples. Our study fills this particular lacuna by evaluating immune cells and secreted factors in the early AMD patient eyes. Much of our current understanding of immune dysregulation in AMD focuses on the damaging, chronic inflammatory processes, with limited understanding of the immunoregulatory or immune dampening mechanisms in the eye. Therefore, we have attempted to address this gap in knowledge in human AMD eyes. Our data not only recapitulates the known elevation of neutrophils and their products, but also identifies significantly elevated, activated NK cells as a key modulator of the innate immune milieu in early AMD eyes, both within the retina and in the intraocular chambers. We report similar findings in an established model of dry AMD ^40,41^. Therefore, our data provides a glimpse of the intraocular immune imbalance in dry AMD eyes that can be a key driver of the disease progression. Importantly, our experimental data demonstrate that replenishing adenosine can reverse the RPE pathology in disease models, suggesting that restoration of intraocular immune balance is a viable, novel strategy for AMD treatment.

## RESULTS

### Immune cell landscape of aqueous humor in early AMD subjects, dry AMD human donor retina and in a mouse model with an atrophic/dry AMD-like phenotype

Study subjects were clinically stratified according to AREDS classification as controls and early AMD (AREDS 2/3) based on retinal imaging wherein we observed hyporeflective sub-RPE deposits at the macular region indicative of drusen **(Figure 1A).** The proportions of NK cells and their functional subsets, along with other immune cell types in the aqueous humor of study subjects and donor retina, were determined by flow cytometry **(Figure 1B; Supplemental Figure 1, 2, 3).** We observed higher proportions of total NK cells, CD16^-^CD56^dim^ NK cells and CD16^-^CD56^bright^ NK cells in early AMD eyes compared to controls, while total leukocytes and CD16^+^CD56^dim^ NK cell proportions remained unaltered between the groups **(Figure 1C-G)**. In-addition, the proportion of NK cell subsets with different activation and inhibitory receptors were altered, particularly activated NK cell proportions, such as NKp44+ NK cells, and NKG2C+ NK cells, were significantly higher in aqueous humor of AMD compared to controls **(Figure 1H)**. The cell surface expression level of individual activating and inhibitory receptors in different NK cell subsets in control and early AMD was observed to be similar based on dimensionality reduction analysis (tSNE-CUDA) **(Supplemental Figure 4A)**. In addition, ten unique cell metaclusters were identified with FlowSOM unsupervised clustering, of which clusters 1, 2, 6 and 8 were CD56+ **(Supplemental Figure 4B).** The phenotypes identified in these metaclusters were CD56^+^NKG2D^+^CD161^+^, CD56^+^CD161^+^CD159a^+^, CD56^+^NKG2C^+^PanKIR2D^+^CD161^+^, CD56^dim^CD16^-^NKG2D^+^CD161^+^ and CD56^bright^CD16^-^NKG2D^+^CD159a^+^ NK cells **(Supplemental Figure 4C)**. Other cells of the innate immune system, such as the classical monocytes and neutrophils, were higher in AMD while intermediate monocytes, non-classical monocytes, and eosinophils remained unchanged **(Supplemental Figure 4D-H).**

**Figure 1.**
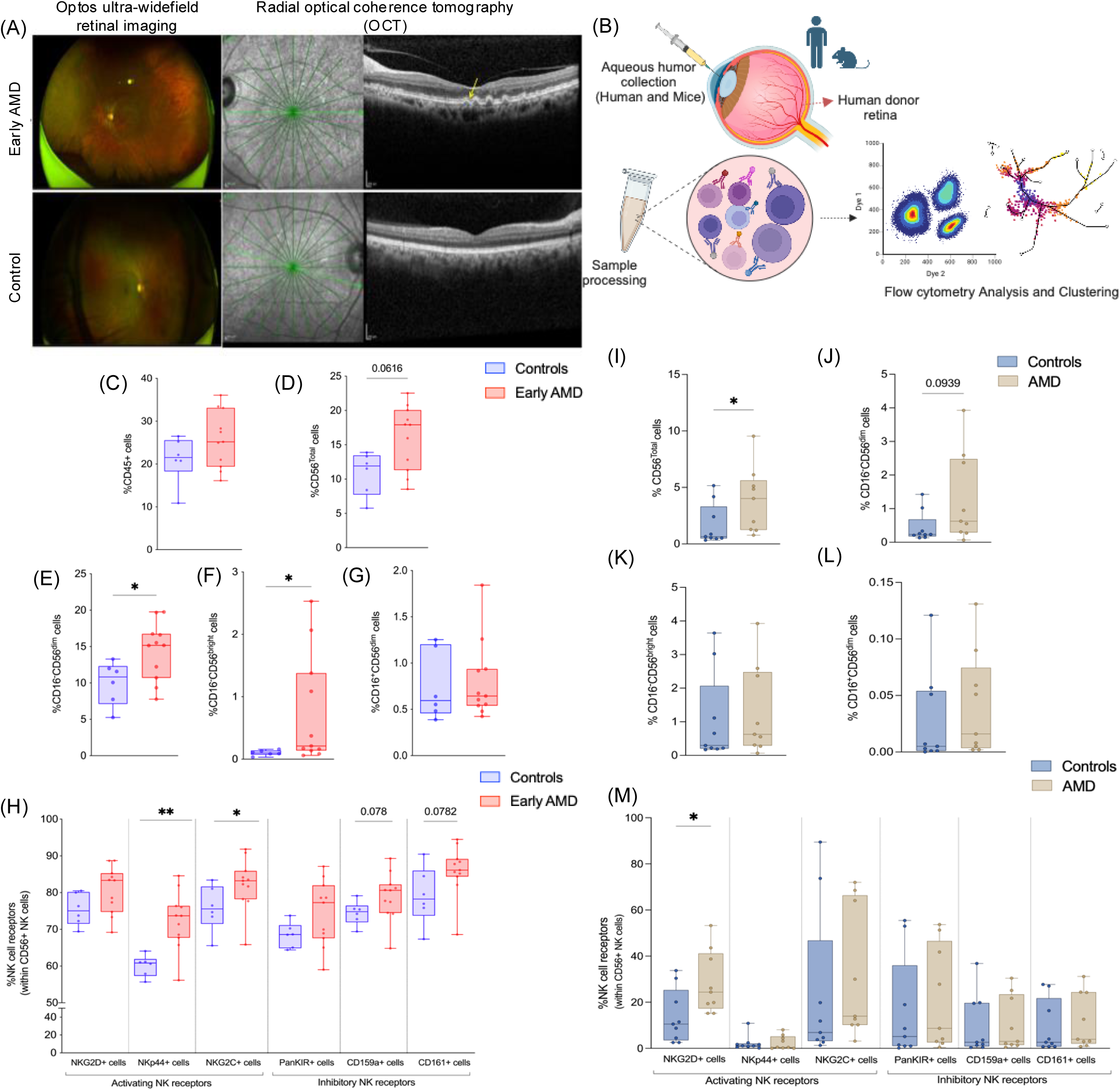
Increase in NK cells and their subsets (with activating and inhibitory receptors) in early AMD subjects and AMD donor retinae. (A) Representative images (top panel left to right) on Optos Ultra-widefield retinal imaging system showed hypopigmented spots at posterior pole and Radial optical coherence tomography (OCT) scan passing through lesion shows hyporeflective sub-RPE deposits (yellow arrow) suggestive of drusen (early AMD) while images in bottom panel (left to right) showed no retinal changes (cataract control). (B) Schema shows aqueous humor and retinal immune cell staining and analysis workflow, created with BioRender.com. Graphs showing increased percentage of (C) CD45+ cells (Leukocytes) (D) CD45^+^CD56^+^ (Total Natural killer cells), (E) CD45^+^CD16^-^CD56^dim+^ (a degranulating NK cell subset) (F) CD45^+^CD16^-^CD56^bright+^ (Cytokine producing Natural killer cells) in early AMD AH, (G) CD45^+^CD16^+^CD56^dim+^ (Cytotoxic Natural killer cells) show no difference in early AMD and control AH. (H) NK cell subsets show increase in proportions of NKG2D^+^ cells, NKp44^+^cells and NKG2C^+^ cells (activating receptors), PanKIR2D^+^ cells, CD159a^+^ cells, CD161^+^ NK cells (inhibitory receptors) in early AMD. Cell proportions were determined by calculating individual counts within total cell numbers acquired. Each data point represented a pool of 5 subjects, 6 data points (cataract control, N=30 subjects) and 11 data points (early AMD, N= 55 subjects). Graph shows increased proportions of (I) CD45^+^CD56^+^ (Total Natural killer cells) and (J) CD45^+^CD16^-^CD56^dim+^(a degranulating NK cell subset); no change in proportions of (K) CD45^+^CD16^-^CD56^bright+^ (Cytokine producing Natural killer cells) and (L) CD45^+^CD16^+^CD56^dim+^ (Cytotoxic Natural killer cells) observed in donor retina of control (n=9) and AMD (n=9). (M) Subsets of NK cells show increased proportions of NKG2D^+^ cells (significant difference seen); NKp44^+^cells, NKG2C^+^ cells (activating receptors) and PanKIR2D^+^ cells, CD159a^+^ cells, CD161^+^ NK cells (inhibitory receptors) show no differences in control and AMD donor retinae. Cell proportions were determined by calculating individual counts within total cell numbers acquired. Box and whiskers plot show Min to Max (all points); *P< 0.05, **P<0.01, Mann–Whitney test.

The immunotypes observed in the aqueous humor was validated in the retina of humor donor eyes with and without dry AMD. The donor eyes for the current study were categorized based on histology assessments as healthy controls and dry AMD. In the human donor retinae, proportions of NK cells and their functional subsets, as well as myeloid cells were determined using flow cytometry (Supplemental Figures 1 and 2). Similar to aqueous humor based findings, dry AMD donor eyes showed increased proportions of total NK cells compared to controls while favorable trend was observed in proportions of CD16^-^CD56^dim^ NK cells **(Figure 1I,J)**. However, proportions of CD16^-^CD56^bright^ NK cells and CD16^+^CD56^dim^ NK cell proportions remained unaltered (**Figure 1L)**. In addition, proportion of NK cells expressing activating receptor-NKG2D^+^NK cells was significantly higher in AMD donor retina compared to controls, though the proportions of NK cells expressing other activating and inhibitory receptor did not show significant alterations (**Figure 1M)**. Of the myeloid cells, classical monocyte proportions were higher in dry AMD donor eye with near significance, while neutrophil proportions were higher in AMD with no significance. The proportions of intermediate monocytes, non-classical monocytes, and eosinophils showed no changes compared to control eyes (**Supplemental Figure 5A-E)**.

An additional cohort of controls and early AMD patients were used to validate the diversity of immune cells and soluble factors in aqueous humor with an extended immunophenotyping panel **(Supplemental Figure 6)** and multiplex bead based ELISA. The proportions of leukocytes, NK cells (Total, CD56^dim^ and CD56^bright^), neutrophils (Total, CD66b^dim^ and CD66b^bright^) were significantly increased in early AMD without changes in monocyte proportions (**Figure 2A-H)** Furthermore, B cells (CD19^+^) were significantly higher in early AMD subjects while no change was observed in proportions of T cells or NKT cells **(Supplemental Figure 7A-C)**. Secreted immune-inflammatory factors such as IFNγ, Perforins, sTNF-R1, TGF-β1 levels were significantly higher while IL-1β, IL-17F, IL-21, RANTES, and BDNF levels were significantly lower in the aqueous humor of AMD subjects compared to controls (**Figure 2I, J; Supplemental Table 1)**. In addition, vitreous humor IFNγ levels in AMD donor eyes were higher compared to controls which served as a validation of the aqueous humor findings (**Figure 2K).** The intraocular immune cell diversity alterations observed in human subjects were validated in the aqueous humor from *Cryba1* cKO mice, which develop an AMD-like phenotype **(Supplemental Figure 8)**. Similar to the patient data, significantly higher proportions of NK cells (total, NK1.1^dim^ and NK1.1^bright^) and neutrophils were observed with monocyte proportions showing an increasing trend. Except leukocytes, proportions of NKT cells and myeloid cells were significantly higher in the aqueous of *Cryba1* cKO mice compared to littermate *Cryba1* floxed controls (**Figure 2L-Q**; **Supplemental Figure 9A, B).** Aqueous humor profiling revealed higher levels of Granzyme, IL-12 - factors expressed by NK cells (**Figure 2 R,S)** and LCN2, MPO – proteins expressed by neutrophils **(Supplemental Figure 10A, B)** in *Cryba1* cKO mice relative to littermate controls. Thus, we investigated NK cells and their interactions in the RPE of the *Cryba1* cKO animals further.

**Figure 2.**
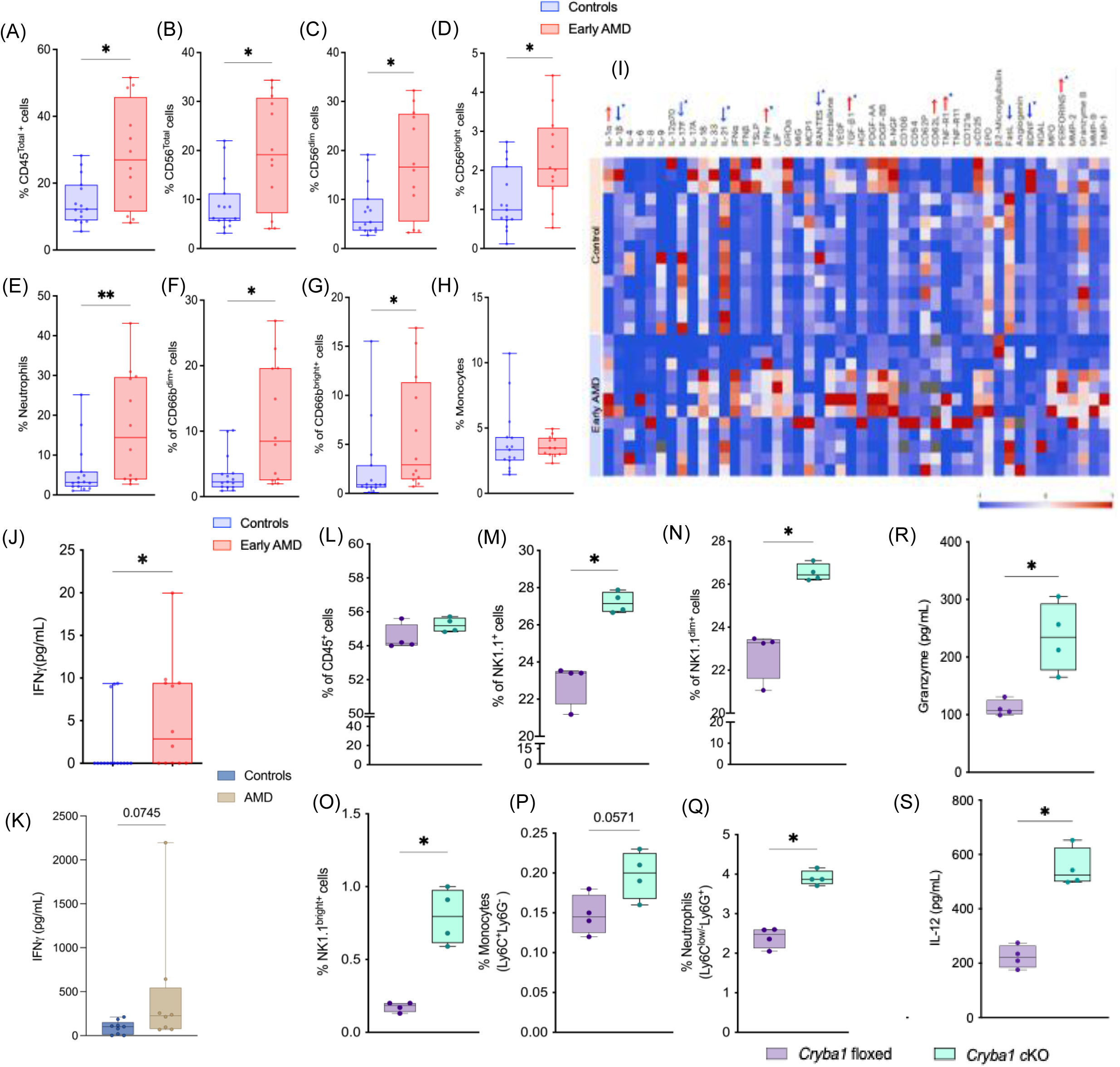
Increased immune cell proportions and elevated levels of NK effector molecules (IFNγ, IL-12, Granzyme) in AH of early AMD subjects, human AMD donor vitreous humor and *Cryba1* cKO mice. (A-H) Graph shows increased percentages of leukocytes, total natural killer cells, cytotoxic NK cells, cytokine producing NK cells, neutrophils (activated and quiescent); monocyte proportions were not altered in AH of individual subjects with controls (n=15) and early AMD (n=12). (I) Heat map showing differential levels of soluble factors in early AMD AH (n=12) compared to controls (n=15). Higher (↑) and lower (↓) soluble factor levels are highlighted, respectively (*significant analytes). (J,K) Absolute levels of IFNγ were significantly higher in early AMD subjects (early AMD, n=12; controls, n=15) and AMD donor eye (AMD, n=8; healthy donor, n=9). (L-Q) Graph shows increased proportions of leukocytes, NK1.1+ NK cells (total, dim and bright), monocytes and neutrophils in pooled AH of *Cryba1* floxed (n=4) and *Cryba1* cKO mice (n=4) at 15 months. (R,S) ELISA showed increased Granzyme and IL-12 levels in *Cryba1* floxed (n=4) and *Cryba1* cKO mice (n=4) at 15 months. Box and whiskers plot show Min to Max (all points); *p value < 0.05, Mann– Whitney test.

### NK cell-related genes and ligand-receptor interactions identified by single cell RNA sequencing (scRNAseq) from the Cryba1 cKO model

Single cell RNA sequencing of the RPE and choroid (cells cleared from choroid blood vessels) of 15-month-old *Cryba1* floxed and *Cryba1* cKO mice showed presence of non-immune and immune cells as shown on UMAP plots using Seurat **(Figure 3A).** Ratio of cell pct showed increased numbers of NK cells, neutrophils and MDSC in *Cryba1* cKO mice RPE. Stacked violin plot represented the expression of marker genes in each cell cluster in mice RPE-choroid complex **(Supplemental Figure 11A,B).** Volcano plot showed genes differentially expressed in NK cell cluster in *Cryba1* cKO mice and littermate controls **(Figure 3B)**. The biological process in Gene Ontology (GO) based enrichment analysis of NK cell population was done in *Cryba1* cKO mice compared to wildtype, where identified genes primarily involved in processes such as regulation of NK cell mediated cytotoxicity, immune responses, and cell killing etc. based on normalized enrichment scores (NES) and were found to be upregulated in *Cryba1* cKO mice **(Figure 3C).** Unidirectional cell-cell communications involving NK cells (denoted as ‘L’, cells expressing ligand) and RPE cells (denoted as ‘R’, cells expressing receptors) were identified by choosing L-R pairs with an interaction score greater than 0.45 (**Figure 3D-F).** The chord plot shows dysregulated L-R interaction pairs as calculated based on the fold changes in interaction scores **(Supplemental Figure 11C)**. The interactions were classified as ‘known’ interactions based on data curated in Cellinker, a tool for understanding cellular networks obtained from peer-reviewed literature on mouse ligand/receptor interaction, while ‘unique’ interactions were those that were not recorded in database. **(Supplemental table 2).** The pathway analysis using KEGG and Reactome showed dysregulated L-R interactions like IFNγ-IFNγR1, Granzyme B (Gzmb)-IGF2R and TNF-LTBR, that are involved in pathways related to programmed cell death including necroptosis, apoptosis and pyroptosis **(Supplemental table 3, 4).** Together these data show the presence of NK cells in mouse RPE along with dysregulated NK cell-RPE interactions that may influence cell survival pathways in the *Cryba1* cKO disease model.

**Figure 3.**
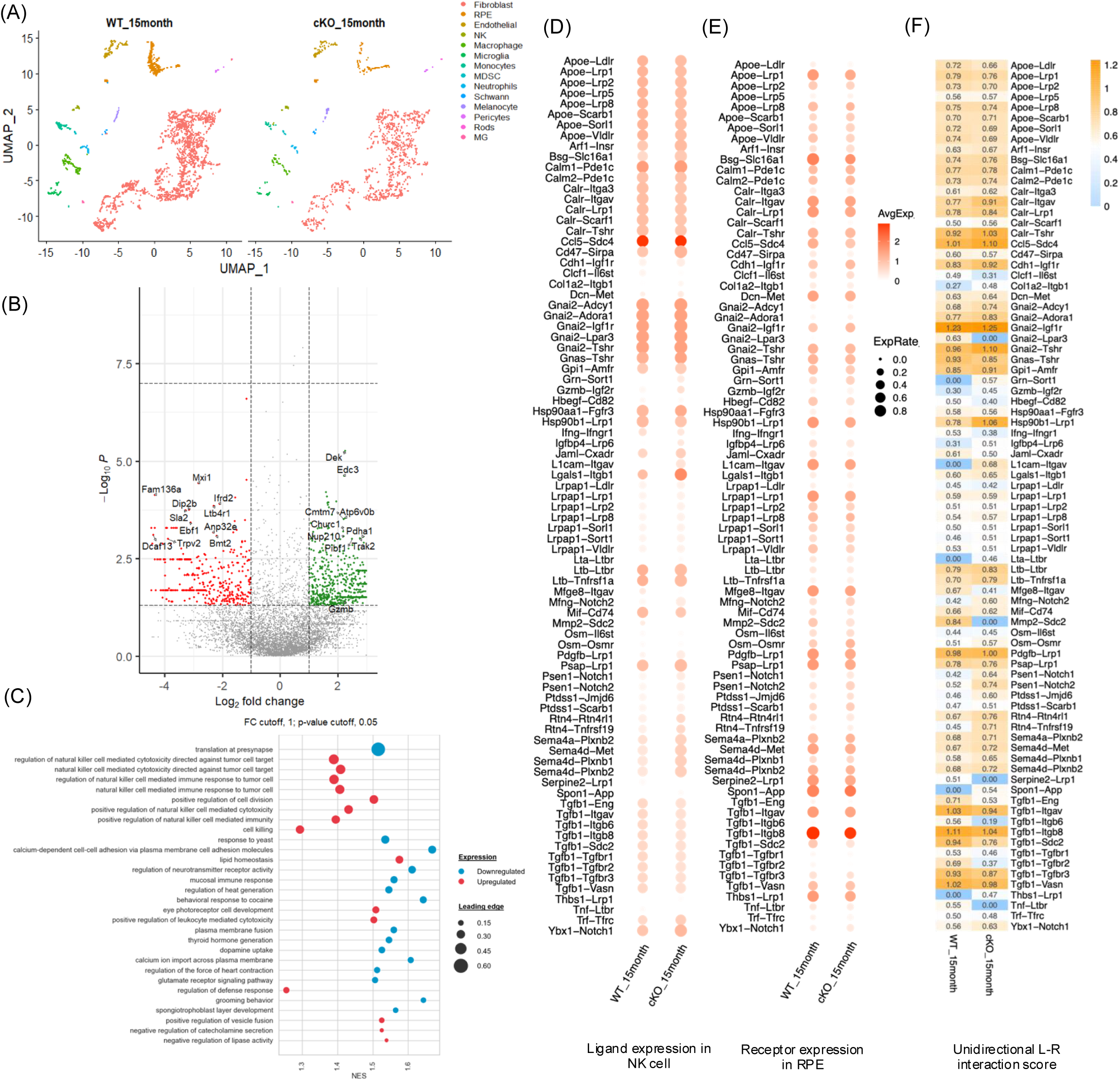
scRNA sequencing of cells from sub-retinal space (SRS: RPE-choroid) in cKO mice and control mice at 15 months shows NK cell clusters and interaction patterns between NK-RPE cell. (A) UMAP plot depicting single cell transcriptomes of different cells types including NK cells, neutrophils, MDSCs in *Cryba1* cKO and control mouse RPE, at 15 months (n=3 in each group). (B) Volcano plot shows differentially expressed genes from NK cell cluster in *Cryba1* cKO mice relative to control. (C) Major gene ontology(GO) functions are related to NK cell cytotoxicity and responses in *Cryba1* cKO mice relative to control at 15months. The size and color of the bubble represents the percentage of identifiers within the biological process and number of genes enriched in the biological process that is of statistical significance, respectively. The process with upregulated genes are indicated in ‘red’ and downregulated genes are indicated in ‘blue’. NES, normalized enrichment score. (D,E) Dot plot comparing ligand expression in NK cells and receptor expression in RPE cells in control and *Cryba1* cKO mice. (F) Ligand-receptor(L-R) interaction scores identified unidirectional NK cell-RPE interaction pairs in control and *Cryba1* cKO mice at 15 months.

### Reduced intraocular immune dampening efficiency in early AMD aqueous humor and AMD donor eyes

Myeloid derived suppressor cells (MDSCs) are immunoregulatory cells of the innate immune system ^42^. The proportion of aqueous humor monocytic MDSC (M-MDSC) cells was lower in early AMD samples compared to controls **(Figure 4A)**. Untargeted metabolomic profiling of patient aqueous humor revealed distinct clustering of control and AMD metabolite profiles **(Supplemental Figure 12A)**. Among the metabolites, an MDSC associated metabolite-adenosine was observed to be markedly lower in early AMD aqueous humor (**Figure 4B; Supplemental Table 5)**. The regulatory relationship between immune factors/cells and immune activation, including NK cells and secreted factors in the aqueous humor, was determined by correlation analysis. The proportion of NK cell subtypes, including NKG2D^+^ NK cells (r = −0.49, *p value:*<0.05), NKp44^+^ NK cells (r = −0.52, *p value:*<0.03), CD161+ NK cells (r = −0.51, *p value:*<0.04) and NKG2C^+^ NK cells (r = −0.51, *p value:*<0.04), were negatively associated with M-MDSC cells. NK cells (total NK, CD56^dim^ and CD56^bright^ cells) were positively correlated with their effector molecules, IFNγ and Perforins. Futhermore, adenosine, a known immunoregulatory factor and NK cell dampener, negatively correlated (*r =* −0.52, *p value:* 0.037) with IFNγ, an immuno-stimulatory factor expressed by activated NK cells **(Figure 4C)**. We quantified M-MDSCs in AMD donor retina to find that compared to control, their proportions were significantly lower **(Figure 4D)**. Furthermore, in retinal lysates from donor retinae, assayed by untargeted metabolomics, PLS-DA score plot showed overlapping patterns in control and AMD with significant reduction in adenosine observed in AMD compared to controls **(Figure 4E, Supplemental Figure 12B, Supplemental Table 6)**. The lower adenosine levels in the aqueous humor as well as retinal lysates from donor eyes was validated by a fluorometric assay (**Supplemental Figure 12C, D)**. Similarly, adenosine was lower in retinal tissue isolated from *Cryba1* cKO mice compared to *Cryba1* floxed littermate controls (**Figure 4F**, **Supplemental Table 7)**, thus corroborating with the observation in patients’ and donor eyes. This observation indicates that homeostatic intraocular immunosuppressive factors are reduced in early AMD.

**Figure 4.**
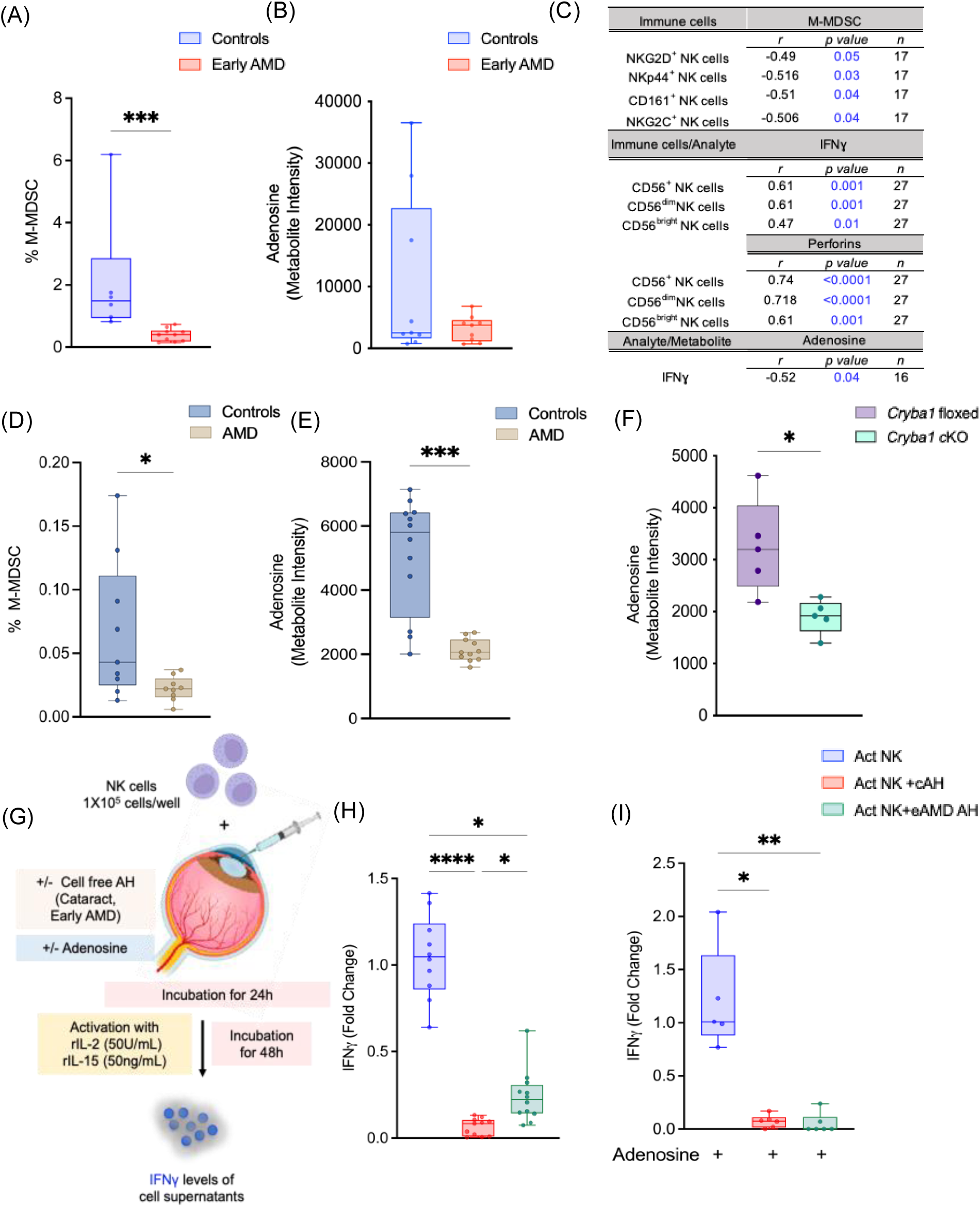
Lower MDSC proportions and adenosine levels (NK dampener) in AMD patient samples with loss of immune dampening function upon addition of early AMD AH. (A) M-MDSC proportions show significant reduction in early AMD AH. (B). Untargeted metabolomics shows lower intensity of adenosine in early AMD AH (n=3) compared to controls (n=3) (metabolomics run in triplicate for each sample). (C) Panel indicates negative correlation between NK cell subsets and M-MDSC in AH (n=17). NK cell subtypes show positive association with IFNγ, perforins, cataract controls and early AMD subjects (n=27); IFNγ shows negative correlation with adenosine (n=16).(D) Reduced M-MDSC proportions in retinal lysates in AMD donor retina compared to healthy retina (n=9 in both groups) (E) Graph shows lower intensity of adenosine in donor AMD retinal lysate (n=4) compared to controls (n=4) by untargeted metabolomics (metabolomics run in triplicate for each sample). (F) Significant reduction in adenosine levels seen in *Cryba1* cKO mice. (G) Method schema showing *in-vitro* assay for assessing immune dampening function in activated NK cells. (H) Graphs show loss of immune dampening due to increased IFNγ levels (represented as fold change) in activated NK cells (isolated from peripheral blood of 5 healthy donors) on addition of early AMD AH versus control AH. (I) Graph shows reduced IFNγ levels (represented as fold change) in activated NK cells (isolated from peripheral blood of 3 healthy donors) on addition of adenosine (5µM). Box and whiskers plot show Min to Max(all points); *P< 0.05, **P<0.01, ****P<0.0001, Mann–Whitney test (A,B,D), Unpaired t test with Welch’s correction (E,F). Kruskal-Wallis test followed by Dunn multiple comparison test (H,I). Table shows r = Spearman rank correlation coefficient, p value<0.05 (C)

Next, the homeostatic intraocular immune dampening capacity of aqueous humor from control and early AMD subjects was determined *ex vivo* by measuring ability of aqueous humor from control and AMD patients in suppressing IFNγ production by activated NK cells **(Figure 4G).** While aqueous humor of both control and early AMD subjects suppressed IFNγ production by activated NK cells, the suppression of IFNγ production by activated NK cells was significantly less in the early AMD group **(Figure 4H)**. This impaired suppression in early AMD aqueous humor was mitigated by addition of adenosine (**Figure 4I)**. These results suggest that the dysregulated intraocular immunosuppressive status in early AMD can be therapeutically rescued.

### Activated NK cells alter RPE barrier function and survival

Since scRNA-seq based analysis including L-R interactions suggested dysregulation in the pathways related to programmed cell death, it suggests a plausible role of NK cells in RPE fate and function. We investigated the role of NK cells in modulating iPSC derived RPE (iRPE) functionality by measuring transepithelial electrical resistance (TEER) ^43^ and the expression of the tight junction protein Zonula Occludens-1 (ZO-1) **(Figure 5A)**. Resistance was measured across iRPE layer co-cultured with activated NK cells (with or without adenosine) at 3 NK cells:1 iRPE cell on matrigel-coated Transwell inserts. Net TEER, a measure of junction integrity, ranged from 160-200Ωcm^2^ at baseline. On addition of NK cells, TEER significantly dropped to less than 80Ωcm^2^ and improved to 110-135Ωcm^2^ with addition of adenosine **(Figure 5B)**. Furthermore, notable disruption in ZO-1 expression was observed in iRPE cultured with activated NK cells, while normal distribution of ZO-1 was observed in cultures where adenosine was also present **(Figure 5C)**. We measured the relative gene expression of IL-1β and secreted levels of IFNγ in iRPE cultures co-incubated with activated NK cells and adenosine. Significant upregulation in mRNA expression of IL-1β, a marker associated with pyroptosis was observed in iRPE cells cultured with activated NK cells which reduced with adenosine **(Figure 5D)**. Similarly, IFNγ levels were measured in cell supernatants of iRPE-NK coculture. Significantly higher secreted levels were seen in cultures with activated NK cells while a reduction was seen in cultures where activated NK cells were treated with adenosine **(Figure 5E)**.

**Figure 5.**
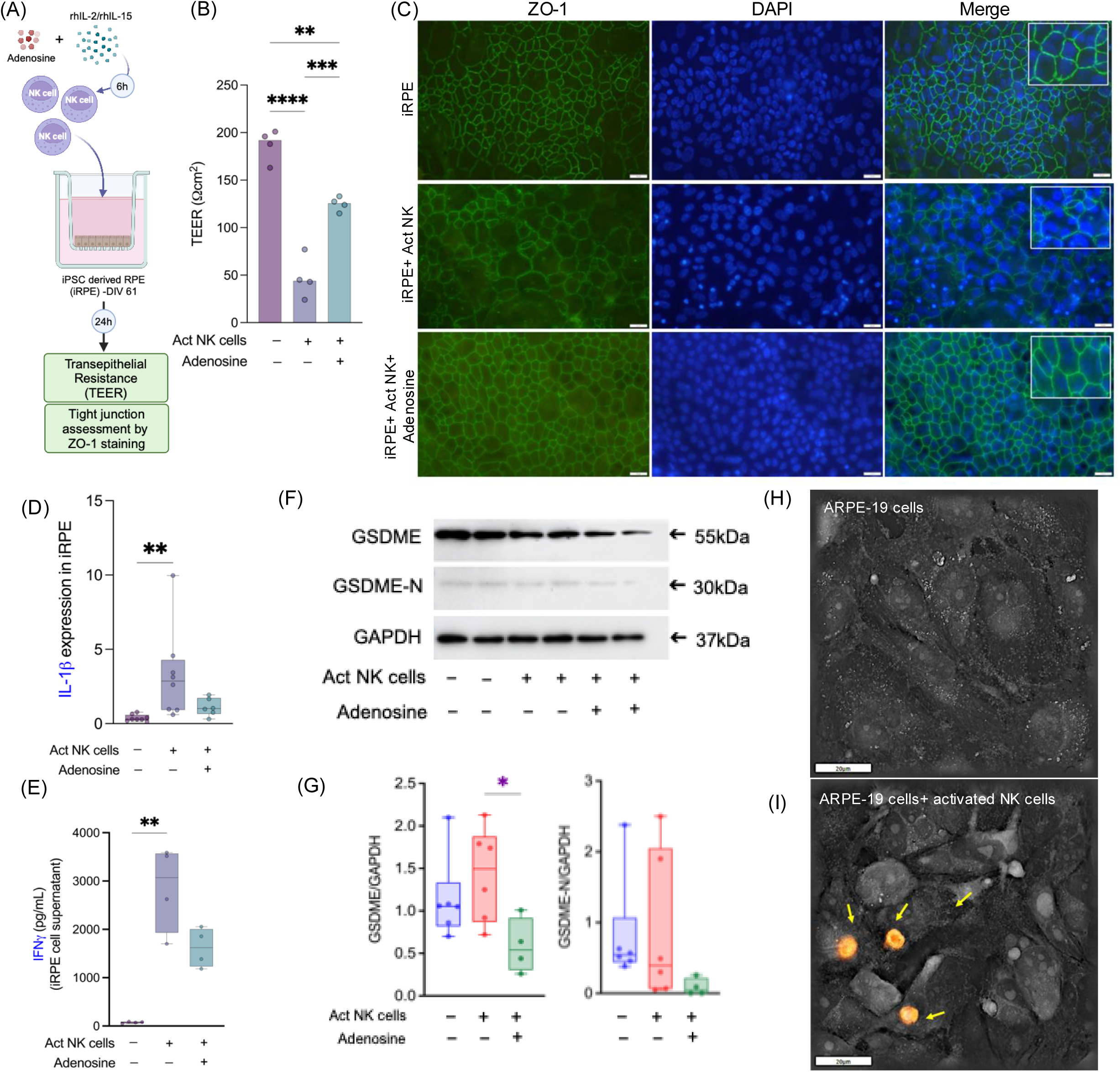
Interaction of RPE with NK cells impairs barrier function and triggers pyroptotic death in RPE. (A) Schema showing procedure to assess effect of activated NK cells and adenosine on iRPE cultured in matrigel-coated Transwell inserts. (B) Net TEER (Ωcm2) of iRPE cells (+/-NK cells and adenosine) shows significant reduction in presence of activated NK cells with significant rescue by addition of adenosine. (C) Zonula occludin (ZO-1) staining of iRPE cells (control cells), iRPE with activated NK cells, iRPE cells with adenosine (5μM) and activated NK cells at 40X magnification; ZO-1-Alexa Fluor 488-Green, DAPI – nuclear stain(blue) and Merge indicated as Blue-Green, in Olympus CKX53 microscope, Scale bar: 20 μm. In presence of activated NK cells, ZO-1 staining and arrangement was disrupted, while with further addition of adenosine the cell morphology was similar to control iRPE cells. Images shown are representative of images obtained on addition of NK cells from 2 donors. (D) Graph shows significant increase in IL-1β mRNA levels, an indicator of pyroptotic cell death, in co-cultures of iRPE cells and activated NK cells; IL-1β mRNA levels are lower on addition of activated NK cells treated with adenosine. (E) Secreted IFNγ levels were higher in iRPE cell supernatants from cells exposed to activated NK cells and reduced after adenosine treatment (5μM) (F,G) Western blot and densitometry of ARPE-19 cells co-cultured with activated NK cells (+/-adenosine) shows increased expression of GSDME in cultures with activated NK cells, with significant reduction on addition of adenosine. Major difference in GSDME-N expression was not seen with activated NK cells, on addition of adenosine, reduction in GSDME-N expression was observed. (H,I) Live cell imaging of NK cell-RPE co-culture was performed on 3D Cell Explorer, NANOLIVE^TM^ for 16h with and without activated NK cells, Scale bar =20 μm. NK cells were stained with PKH26 Red Fluorescent cell membrane cell linker kit (orange colored).Figure H shows no changes in morphology/vacuolations were observed in ARPE-19 cells. Figure I shows activated NK cells interacting with ARPE-19 cells suggestive of pyroptosis like increased cell swelling and localized vacuolations (vacuolations on ARPE-19 cells are indicated with yellow arrows). Graphs show Min to Max (all points); P values (< 0.05) obtained using Ordinary one way ANOVA followed by Tukey’s multiple comparison test.

We also investigated the potential additive effect of IFNγ in NK cell mediated RPE dysfunction using polarized ARPE-19 cells co-cultured with activated NK cells (5 NK cells:1 ARPE-19 cell) in presence/absence of IFNγ (50 ng/mL) **(Supplemental Figure 13A).** We observed a reduction of TEER ^44–46^ in polarized ARPE-19 cultured with activated NK cells which further reduced in presence of IFNγ **(Supplemental Figure 13B)**. Similarly, disruption in ZO-1 staining was observed in polarized ARPE-19 cells treated with activated NK cells (+/-IFNγ) **(Supplemental Figure 13C)**. Image J based quantification with TightJunction analysis plugin measured cell-cell edge quantification parameters like tight junction length, Euclidean distance, and edge ratio. Significantly altered ZO-1 organization was calculated in polarized ARPE-19 cells co-cultured with activated NK cells based on total length and mean area. We observed significant reduction in total length and significant increase in mean area in polarized cells co-cultured with activated NK cells (+/-IFNγ) **(Supplemental Figure 13D, E)**. These observations confirm that activated NK cells can dysregulate RPE function. LDH activity assay demonstrated NK cell mediated cytotoxicity in ARPE-19 cells with increase in Effector:Target ratios (E:T) irrespective of IFNγ treatment, where at ratios of 3:1, 6:1 and 9:1 about 24%, 33% and 43% cytotoxicity was observed; respectively **(Supplemental Figure 13F)**. Furthermore, with Annexin V-7AAD staining, the proportions of early and late apoptotic cells were higher with increased NK to ARPE-19 cell ratio **(Supplemental Figure 13G-K**).

Activated NK cells release factors like IFNγ and cytotoxic granules like perforins and granzyme B. Perforins help in the entry of Granzyme B into cells, thereby activating its signaling pathway and regulating expression of Gasdermin-E (GSDME) which forms pores that rupture the plasma membrane and release cellular factors ^47^. Here, expression of GSDME showed an increasing trend in ARPE-19 cells co-cultured with activated NK cells, while GSDME and GSDME-N reduced on incubation with adenosine (5μM) treated activated NK cells **(Figure 5F, G)**. RPE cell membrane changes were seen with live cell imaging upon interaction with activated NK cells in contact-dependent co-culture model. These RPE cells developed localized vacuolation (membrane pores) and appeared swollen, an indicator of pyroptotic cell death. Control cells showed no changes in morphology nor did they exhibit vacuolations **(Figure 5H, I; Movies 1,2)**. Cytoplasmic density of ARPE-19 cells was increased after interacting with activated NK cells compared to control cells, as seen on 3D Refractive index maps **(Supplemental Figure 14A, B)**. Collectively, these observations suggest pyroptosis may be another mechanism by which NK cells induce RPE cell death. Therefore, our findings suggest that the NK cell – RPE cell axis detrimentally impacts RPE function and survival.

### Adenosine treatment lowers aqueous humor immune cells and Gasdermin E-mediated pyroptotic cell death in RPE

Adenosine is a known suppressor of cellular immune responses and can modulate immune cells to switch their functional status ^48–50^. Its role in suppressing NK cell functional responses, cytolytic activity and its inhibition of NK cell granule exocytosis have been reported in other models ^51–53^. To test the therapeutic potential for NK cell suppression *in vivo*, we administered adenosine (4μM) intravitreally in one eye while PBS was injected in the contralateral eye of 10 month old *Cryba1* cKO mice. We observed a significant reduction in aqueous humor levels of adenosine in *Cryba1* cKO while significantly higher adenosine was observed in adenosine-injected *Cryba1* cKO eyes relative to contralateral vehicle-injected eyes **(Supplemental Figure 15).** Interestingly, adenosine-treated *Cryba1* cKO mice showed significantly lower proportions of NK cells (total, NK1.1^dim^, NK1.1^bright^), monocytes and neutrophils in aqueous humor **(Figure 6A-E**). Leukocytes and CD11b^+^ myeloid cells did not show significant difference in proportions while proportions of Natural killer T cells were significantly higher in *Cryba1* cKO and significantly reduced in adenosine-injected *Cryba1* cKO compared to *Cryba1* cKO mice (**Supplemental Figure 16A-C)**. Significantly lower levels of Granzyme in adenosine-injected eyes of *Cryba1* cKO mice validates the function of adenosine in inhibiting exocytosis of NK cell lytic granules **(Figure 6F)**. To validate presence of granzyme in eyes from dry AMD, we prepared paraffin embedded sections of normal and AMD donor retina. We observed increased staining of granzyme B (GzmB) in AMD donor eyes compared to control (**Figure 6G).** H&E staining of AMD donor eyes also demonstrated characteristic drusen deposits and gliosis indicative of dry AMD phenotype, while no retinal pathologies were observed in control eyes (F**igure 6H**). We also measured GSDME and GSDME-N in RPE lysates of *Cryba1* floxed and *Cryba1* cKO mice with and without adenosine injection. Levels of GSDME and GSDME-N in RPE lysates was higher in the *Cryba1* cKO eyes compared to *Cryba1* floxed mice and was reduced upon adenosine-injected eye **(Figure 6I-K).** These results suggest that targeting NK cells with adenosine to inhibit NK cell mediated pyroptotic cell death is a potential therapeutic approach for early AMD management.

**Figure 6.**
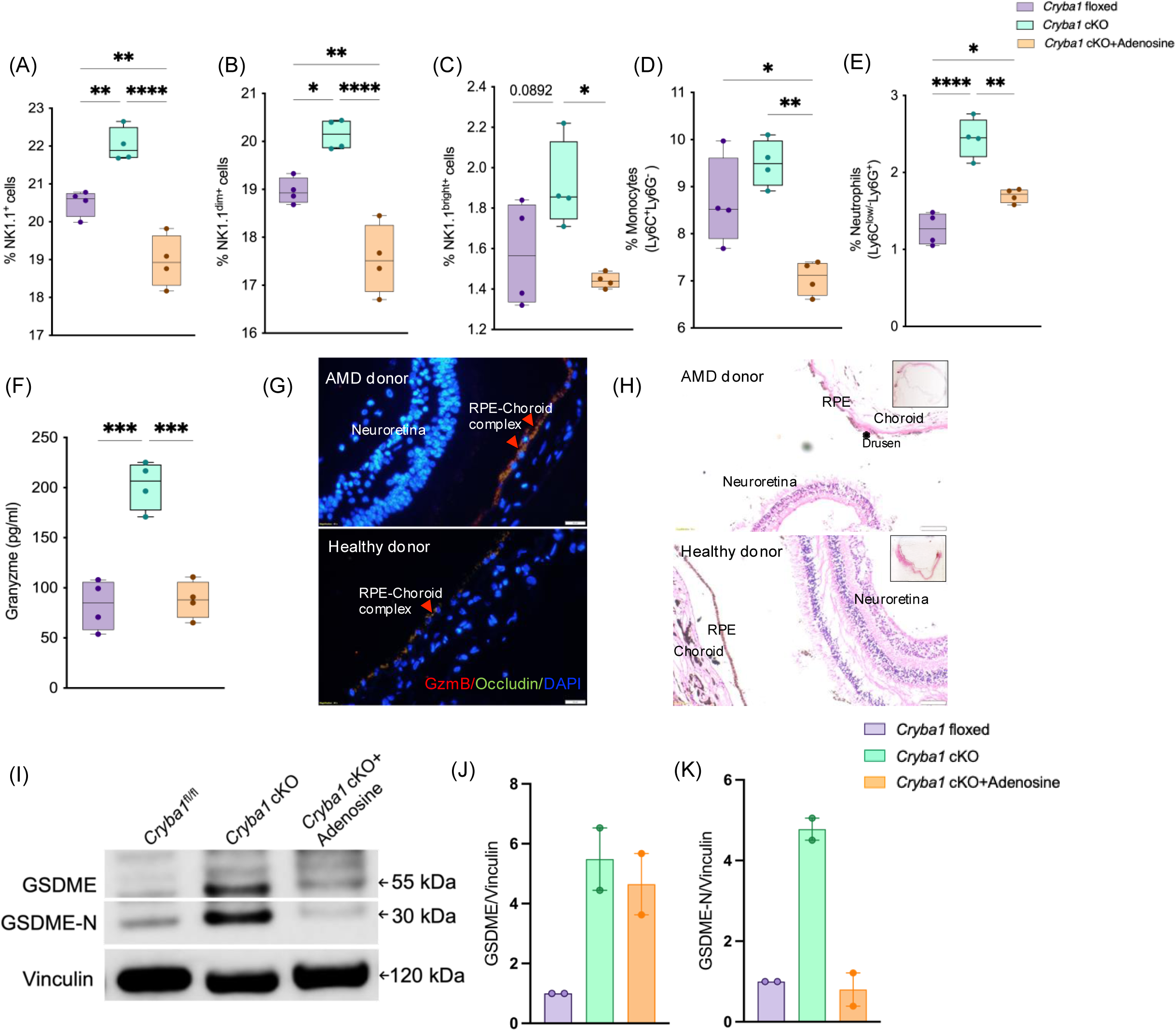
Intravitreal adenosine reduces aqueous humor immune cell infiltration and rescues NK cell-RPE interaction dependent pyroptotic death *in-vivo*. (A-E) Lower proportions of NK1.1+ NK cells (total, dim and bright), monocytes and neutrophils found in pooled AH of *Cryba1* cKO mice injected with adenosine (n=4) compared *Cryba1* cKO mice prior to intravitreal adenosine injection. (F) Granzyme levels as measured by ELISA are elevated in AH from 15 month old *Cryba*1 cKO mice, but reduced to level of control mice post intravitreal adenosine injection (n=4). (G) Representative immunofluorescence images (40X magnification) of human donor retina and RPE choroid complex stained with Granzyme B and Occludin antibodies, Scale bar: 20µm (n=3 each). (H) Representative phase contrast image (10X magnification) showing H&E-stained tissue section of AMD show characteristic drusen deposition (indicated by ∗) in RPE layer, no retinal changes are seen in healthy donor (control) eye (n=3 each), Scale bar: 100µm. Inset shows H& E stained eye globe slice in AMD and healthy donor respectively (I-K) RPE cell lysates show increasing trend in expression of GSDME and GSDME-N in *Cryba1* cKO mice and reduces in RPE lysates of adenosine treated eyes. Box and whiskers plot show Min to Max(all points), *P< 0.05, **P<0.01,****P<0.0001, Ordinary one-way ANOVA followed by Tukey’s multiple comparison test (B-H, J, K).

## DISCUSSION

Age-related diseases, including dry AMD, are characterized by chronic inflammatory processes that directly contribute to tissue degeneration ^54^. Immunologic triggers such as lipids, complement factors, APOE, albumin, amyloid-β, advanced glycation end products, etc., that accumulate in the RPE and are inadequately cleared by compromised autophagy have been demonstrated in AMD models ^55^. In patients, RPE changes are clinically noted early in disease ^56^. Cumulatively, these triggers activate retinal microglia, release of complement components, inflammasome activation, and secretion of growth factors like VEGF and pro-inflammatory factors ^57^. This cascade recruits a variety of immune cells, including monocytes, neutrophils, macrophages, T cells, and mast cells, which create a persistent intraocular inflammatory microenvironment in AMD models ^58^. Currently, targeted therapies exist only for GA (targeting complement factors C3), and wet AMD (including photocoagulation, intravitreal anti-VEGF, and steroids), leaving the largest AMD population, those with early and intermediate AMD, without specific treatment options ^59^. Thus, a strategy that can restore homeostasis by reversing the immune imbalance in the early stage of disease has potential not only for preventing disease progression, but also disease remission. Therefore, it is critical to understand the intraocular milieu in such early-stage patients to improve our understanding of the disease, but also to identify potential avenues for therapies that can prevent further disease progression.

RPE degeneration, a primary pathological driver^5^ of AMD and a clinically observable indicator of disease progression, has been linked to dysregulation of the intraocular immune milieu. In prior investigations involving animal models exhibiting AMD-like characteristics, infiltrating neutrophils, neutrophil traps, and subsequent microglial activation have been shown to contribute to retinal disease progression ^18^. Interestingly, we found a concurrent reduction in the M-MDSC cell proportions as well as reduced levels of the immune-dampening metabolite, adenosine, in the dry AMD patient eyes. This finding is mechanistically important for defining how innate immunity becomes imbalanced in early stages of disease. NK cells play a pivotal role in the innate immune defenses by exerting cytolytic activity against stressed cells, pathogens, and tumor cells ^60^. This activity is regulated by a repertoire of inhibitory and activation receptors on the cell surface, which modulate their effector functions as has been demonstrated in obesity ^61^, cancers ^62^, infectious diseases including COVID-19 ^63^, and degenerative diseases like AD and Parkinson’s disease (PD) ^27^. In support of our observations regarding NK cells, examination of published scRNA sequencing data from AMD eyes (human and CNV model of mice) shows NK cell signatures ^64, 65^. Notably, activated NK cells secrete factors such as IFNγ and cytolytic granules like Perforins and Granzyme B, which consequently induce cytotoxicity in the interacting cellular layers. While such activity is desired for tumor cell clearance ^66^, an aberrant NK cell activation can potentially damage the RPE, as it does to neurons in PD ^67^. In our study, elevated levels of IFNγ and Perforins, secreted factors that have NK effector activities, were present in dry AMD eyes, suggesting a mechanistic link between aberrantly activated NK cells and RPE changes.

The role of MDSCs in AMD is particularly intriguing, as these cells have been implicated in suppressing aggravated immune responses ^33^, though their levels in human dry AMD eyes had not previously been assessed. MDSCs express CD39/CD73 ectoenzymes that convert ATP to extracellular adenosine ^68^, which serves as a crucial immune modulator. In the retina, CD39 is expressed in the optic nerve head, retinal vasculature, and microglia, while CD73 is found in the photoreceptor layer ^69^. This distribution helps maintain optimal adenosine levels within the intraocular milieu. Adenosine-dependent immunomodulatory signaling has roles in anti-inflammatory responses, lysosomal clearance, and cell survival ^70^. In tumor tissues, elevated adenosine secretion by MDSCs hinders NK cell surveillance, reduces cytotoxic activity through the perforin/granzyme B or FasL-dependent cell death pathways, and restricts the release of IFNγ ^52,71^. Unlike in the tumor microenvironment, we observed reduced proportions of MDSCs and lower levels of FasL, adenosine in the aqueous humor of AMD patients, indicating a deficiency of immune dampeners in the AMD ocular environment. Importantly, we demonstrate that the reduced immune dampening capacity of AMD AH, compared to non-disease AH, compromises its ability to restrict NK cell mediated IFNγ secretion and that this effect can be rescued by treatment with adenosine. Therefore, chronic, aberrant NK cell activation coinciding with failure to suppress NK cell function in early AMD can be an important cause of RPE dysfunction and death.

In various ocular conditions, such as diabetic retinopathy and glaucoma, a novel inflammatory programmed cell death pathway known as PANoptosis has been proposed ^72^. PANoptosis involves the coordinated activation of multiple cell death pathways, including apoptosis, necroptosis, and pyroptosis. Unlike the canonical inflammasome mediated pathways, pyroptosis may be facilitated via Granzyme/Perforin complexes driving Gasdermin mediated cell death as demonstrated in cancer models^73^ and ocular surface cells^74^. We found higher expression of Granzyme B in the donor AMD RPE cells compared to non-disease controls, suggesting that infiltrating NK cell mediated pyroptosis is directly linked to RPE dysfunction. In co-culture models, we show that NK cells can induce RPE dysfunction and initiate pyroptosis-driven cell death in RPE cell monolayers through the activated NK cell IFNγ/Perforin/Granzyme B pathway. Importantly, supplementing adenosine in these assays, reversed the NK mediated effects on RPE, which were recapitulated *in vivo*, in the AMD disease model. The data therefore suggests that exogenously administered adenosine, can dampen NK cell function, which may consequently prevent RPE cell dysfunction and death in dry AMD patients.

In conclusion, our study reveals a new functional role for NK cells, encompassing both cytotoxic and cytokine-producing subtypes, as well as key effector factors such as IFNγ and Perforins in AMD pathology. Importantly, the loss of MDSCs and lower adenosine levels negatively correlated with NK cell numbers and IFNγ levels, a new observation in early AMD. This skewed inflammatory milieu driven by activated NK cells can initiate the Perforins/Granzyme B pathway for the elimination of impaired RPE cells in a Gasdermin E-dependent manner. Taken together, our findings contribute to a deeper understanding of the immunological mechanisms operational in AMD and offer new avenues for therapeutic interventions using a biological modality.

## METHODS

### Study cohort and design

A cross-sectional study was approved by institutional ethics committee and performed as per the guidelines of Declaration of Helsinki and subjects were recruited following informed written consent. Age-matched study subjects include patients that underwent cataract surgery and then classified into cases with and without macular changes. Age-related eye disease study (AREDS) defined severity of AMD combining baseline clinical findings like drusen, RPE abnormalities, atrophy and neovascularization. In this study, subjects graded as AREDS 2 (early stage) and 3 (intermediate stage) were grouped as early AMD and AREDS 1 were considered as cataract controls. The following subjects were excluded from the study: (i) those with any clinically significant underlying systemic, autoimmune or other inflammatory conditions; (ii) those with active uveitis, glaucoma; (iii) those with ongoing or history of surgical or medical treatment for retinal conditions like proliferative diabetic retinopathy or retinal vascular diseases. A total of N=112 patients with n=45, cataract control and n= 67, early AMD group was investigated in the entire study where aqueous humor was the sample source. The aqueous humor pooled samples were prepared from 30 subjects (pool of 5 subjects per data point), control, n= 6 data points and from 55 subjects (pool of 5 subjects per data point), early AMD, n= 11 data points. The aqueous humor samples from remaining subjects were run as individual samples for validating the immune cells and identifying dysregulated soluble factors in control, n=15 and early AMD, n=12.

Human donor eyes included in the study was collected from Dr. Rajkumar Eye Bank, Bangalore, India and Shankar Anand Singh Eye Bank, Bangalore, India as per Eye Banking Standards of India 2020 and tenets of the Declaration of Helsinki. Enucleated eyes were dissected and samples collected for the study include vitreous humor; retinal tissue for immunophenotyping, metabolic analysis; and one half of eyeball for preparation of paraffin embedded tissue blocks for histology. Human donor eyes were classified into healthy controls (n=9) and dry AMD (n=9) based on histology.

### Animals

All animal studies were conducted in accordance with the Guide for the Care and Use of Animals (National Academy Press) and were approved by the University of Pittsburgh Animal Care and Use Committee (Protocol # 23104041). The *Cryba1* cKO mice and the litter controls were bred as described previously ^9,75,76^. The animals were housed and maintained by trained veterinarians in the Mercy Pavilion animal facility, UPMC.

### Phenotyping of immune cell subsets in human aqueous humor and donor retina

Aqueous humor sample collection and processing as well as step-wise processing of donor retinae are detailed in Supplemental methods. Immune cell proportions in patient aqueous humor and donor retinae were determined using cell specific fluorochrome conjugated antibodies by flow cytometry. Briefly, for staining the aqueous humor was centrifuged at 400g for 5 minutes at 4°C. In donor retinae, single cell suspensions prepared was centrifuged at 400g for 5 minutes at 4°C. The cell pellet obtained in both cases was stained with antibody cocktail prepared in staining buffer (5% Fetal Bovine Serum in 1X Phosphate Buffer Saline, pH 7.4) for 45 minutes at room temperature with gentle agitation. Cells were washed by centrifugation and resuspended in 300 μL 1X Phosphate Buffer Saline, pH 7.4. Data acquisition for cells isolated from aqueous humor and donor retinae was done on BD FACS Lyric (data set in Fig 1) and BD FACS Canto II flow cytometers (data set in Fig 2) respectively. Data acquisition was done using BD FACSuite (BD FACS Lyric) and BD FACSDiva software (BD FACS Canto II), BD Bioscience, USA followed by data analysis with Kaluza software (Beckman Coulter, USA). The antibody titrations used have been detailed in supplemental methods. Post-acquisition compensation was done using single stained controls and fluorescence minus one controls. Cell populations were identified by manual gating strategy (Supplemental Figure 1, 2, 3 and 6) and regions were represented based on positive staining. Cell proportions were determined based number of cells stained positive for the respective antibody to total cell number acquired for each sample. Cytobank plugin was used for dimensionality reduction and clustering analysis.

### Soluble factor profiling of human aqueous humor, vitreous humour and RPE cell supernatants

The levels of soluble factors in the aqueous humor (CBA, LEGENDplex^TM^), vitreous humour and cell supernatants from iRPE cultures (IFNγ,IL-1β-Legendplex^TM^) were measured by multiplex ELISA. Samples for Cytometric Bead Array (CBA), BD^TM^ CBA Human Soluble Protein Flex Set System (BD Biosciences, USA) and LEGENDplex^TM^ (BioLegend, USA) kits were used as per manufacturer’s instructions. The analytes measured using each kit is elaborated in supplemental methods. Briefly, samples was incubated with capture bead and detection reagent; post incubation the beads were washed and acquired on BD FACSCanto II, BD Biosciences, USA. The signal intensities of each analyte were recorded and the absolute concentration was calculated using analyte specific standards. Data extraction was done using the FCAP array Version 3.0 (BD Biosciences, USA) or LEGENDplex™ Data Analysis Software (BioLegend, USA).

### Untargeted metabolomics of human aqueous humor and donor retina

Sample pre-processing was required for retinal lysates. Briefly, a small portion of retina was immersed in 0.1M Triethylammonium bicarbonate buffer (TEAB). The samples were lysed by sonication (power: 40V, energy: 20%, pulse: 5) and incubation in ice for 1min, three times. The mixture was centrifuged at 13,200rpm for 15min to remove debris and supernatant was further processed.

For untargeted metabolomics, 30μL aqueous humor samples/retinal lysates were subjected to solvent based extraction in 600μL of ethanol/methanol mixture (v/v, 1:1). Samples were centrifuged at 4 °C for 15 mins to remove debris after vigorous vortex. The samples were vacuum dried and pellet was resuspended with 80μL 50% acetonitrile ^77^. Samples were loaded on ExionLC (Ultra High Performance Liquid chromatography UHPLC) coupled to Triple-TOF 5600 (Mass spectrometer), SCIEX, MA, USA. The raw MS data were converted to mzML files and metabolite features were extracted and annotated using a cloud based tool, Metaboquest, Omicscraft, DC, USA. Partial least-square discriminant analysis (PLS-DA) was calculated based on variability between the control and early AMD groups and the variable importance on projection (VIP) was determined. Metabolites with VIP values >0.4 and P-values <0.05 were considered for analysis.

### Adenosine assay by ELISA

Aqueous humor levels of Adenosine (μM) from control/early AMD subjects and retinal lysates from donor retina were measured using Adenosine Assay kit (MET-5090,Cell Biolabs, USA) according to manufacturer’s protocol. The aqueous samples were diluted to 1:7 and retinal lysate prepared by sonication were diluted to 1:5. Briefly, 50μL sample and standards were added to black microtiter plate in pairs where to one set 50μL Reaction mix and to second set 50μL control mix was added respectively. The well contents were mixed well and incubated for 15minutes at room temperature in dark environment. Fluorescence was measured at excitation wavelength 530nm and emission wavelength 590nm on Spectramax.

### Immunophenotyping of mouse aqueous humor by flow cytometry

Samples collected from mice eye were placed on ice and centrifuged at 800 g for 4 minutes. The cell pellet was used for assessing the percentage of neutrophils and NK cells, after incubating in blocking buffer (0.5 % BSA, 2 % each of goat, rat and mouse Serum in PBS) for 45 minutes and then staining with antibody cocktail prepared in Flow Cytometry Staining Buffer(*# 00-4222-26, ThermoFisher, USA) at a concentration of 1 μg/mL for 90 min at room temperature. The pellet was washed with staining buffer thrice and then acquired using the BD LSR II, BD Biosciences, USA). Post-acquisition analysis and compensation was done using single stained controls on Kaluza software, Beckman Coulter, USA. Cell proportions were determined by manual gating strategy **(Supplemental Figure 6)** based number of cells stained positive for the respective antibody.

### Untargeted metabolomics from mouse RPE tissue

The RPE tissues from 4-month-old *Cryba1* cKO and its litter control mice were lysed to perform mass spectrometry analysis for untargeted metabolites as a paid service from the Metabolite Profiling Core Facility at the West Coast Metabolomics Center, UC Davis.

### ELISA based profiling of mouse aqueous humor

The levels of Granzyme (R&D Systems, DY1865-06), IL-12 (Invitrogen, BMS616), LCN2 (Invitrogen, EMLCN2) and MPO (Invitrogen, EMMPO) in the aqueous humor isolated from mice were determined by following the manufacturer’s instructions. The optical density at 450nm was measured with Synergy H1(Biotek, USA) plate reader. The levels of each factor was calculated with respect to their standard curve prepared using respective standards.

### Single cell RNA (scRNA) sequencing of *Cryba1* mouse RPE

Dissected SRS (RPE-choroid) from 15-month-old *Cryba1* floxed and cKO mice was made into single cell suspension post removal of immune cells from choroid blood vessels as described previously^9^. The cell suspension was subjected to scRNA sequencing as a paid service from the Genomics Research Core of University of Pittsburgh to identify the different cell types and transcript profiles of each cell type in mouse RPE. Average expression values of the genes in each cell type and each sample were calculated by the function “Average Expression” in the Seurat package. Differential expression analysis was performed on each cell type of interest between floxed and cKO samples by the function FindMarkers with test.use = “wilcox” in the Seurat package. For the analysis of ligand-receptor interactions between cells, we used the default built-in ligand-receptor interaction database in the package LRLoop^78^. Then we scored the candidate ligand-receptor pairs between each pair of cell types of interest in each sample by the SingleCellSignalR ^79^.,

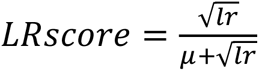

where l is the average expression value of the ligand gene, r is the average expression value of the receptor gene, μ=mean(C) and C is the normalized read count matrix.

### NK cell activity assay

IFNγ release was used as a readout to assess activity of NK cells. Isolated NK cells were activated in culture with rhIL-2 (500U/mL) / rhIL-15 (50ng/mL) cocktail at 37°C for 48h. The influence of control and early AMD aqueous humor with/without adenosine on NK cell activity was determined by incubating NK cells with AH or adenosine for 24h, followed by rhIL-2/ rhIL-15 activation for an additional 48h at 37°C. The cell-free supernatants from these cultures were used for IFNγ measurement using LEGENDplex^TM^, Biolegend, USA according to manufacturer’s instructions.

### iPSC to RPE differentiation

iRPE was derived from commercial NCL1 hiPSC line passage 20, using a modification of a previously described protocol by D.O. Clegg *et.al*, 2017 ^80^. For differentiation to iRPE, iPSC were seeded into Matrigel^R^ Growth Factor Reduced (354320, Corning, USA) coated six-well plates. iPSC colonies formed a confluent

monolayer at 70-80% confluence, transition to retinal differentiation media 1 (RDM1) for three days [DMEM/F12 with antibiotics, supplements and small molecules like N2 supplement (100X, 17502048, Gibco), B27 (50X, 17504044, Gibco), non-essential amino acid (100X, 11140050, Gibco), nicotinamide (10mM, N3376, Sigma aldrich), Noggin (50ng/μL, 120-10c Peprotech), Dkk1 (10ng/μL,#5439DK, R&D System), IGF-1 (20ng/μL, I1146 0.1MG, Sigma aldrich)]. At the end of 3 days, cells were transitioned into RDM2 media [RDM1 media+ B-FGF (5ng/μL,13256029,GIBCO)], followed by RDM3 media [RDM2 media+ Activin A (100ng/μL, A4941,Sigma aldrich)], RDM4 media [RDM3 media+ SU5402 (10μM, s-204308, Chemcruz)] and RDM5 media [RDM4 media+ CHIR99021 (3μM, SML 1046, Sigma)], where cells were cultured in each media for 3 days to ensure RPE committed fate. Immature RPE cells were transitioned into RPE supporting media (RSM media) [DMEM/F12 with antibiotics, non-essential amino acid, B27, Fetal bovine serum-United States (3%, 26140079, Gibco), Sodium pyruvate (100mM, 11360070, Gibco)] for 40 days or more for further assays.

### Barrier function of iRPE cells and polarized ARPE-19 cells

iRPE (DIV115) and ARPE-19 monolayer cultures were established by seeding 2 × 10^5^ cells on 12 well-transwell insert (PET, 0.4 μm pore, 24 mm). On cell attachment, cells were supplemented every 2 days with respective media. Barrier function was assessed by monitoring transepithelial resistance (TEER) using a Millicell ERS-2 Voltohmmeter, Merck Millipore, USA. Resistance (Ωcm^2^) of RPE monolayer was calculated using average of four independent measurements and corrected using background resistance (filter with culture medium alone) to determine barrier integrity. Zonula occludens-1 (ZO-1), a tight junction protein was visualized by immunofluorescence to verify barrier function. Briefly, the cell-bound membrane filters were fixed with 100% methanol, washed, blocked and stained overnight at 4°C with ZO-1 antibody. Membrane was washed and stained with secondary antibody to ZO-1 tagged with Alexa Fluor 488. The membranes were carefully mounted on slides and stained with DAPI. Imaging was done using an Olympus CKX53 microscope at 20X magnification.

### LDH activity assay

ARPE-19 cells (1X10^5 cells) were seeded in 96-well plates for 24h. Act NK cells were co-cultured with ARPE-19 at 3:1, 6:1, 9:1 (effector to target ratio) in presence or absence of rhIFNγ (50ng/mL). Cell death was assessed by LDH activity using Lactate Dehydrogenase Activity Assay Kit (MAK066,Sigma Aldrich, Merck, USA) at the end of 24h according to manufacturer’s instruction. Briefly, 50uL of cell free supernatants was added to 50uL Master reaction mix (48uL LDH Assay buffer+ 2uL LDH substrate mix) and mixed well. Initial reading was taken within 2-3 minutes of addition of reaction mix. Measurements were taken every 5minutes for 30minutes duration. Colorimetric measurement was done at 450nm using Spark Multimode plate reader, Tecan, USA. The measurements were calculated based on NADH standards and LDH activity was calculated as percentage of LDH release.

### Cytotoxicity assay

AnnexinV-7AAD staining was used to determine the fate of ARPE-19 cells following co-culturing with activated NK cells. Briefly, ARPE-19 (40000 cells/well) stained with PKH26 red cell linker membrane labelling dye (MIDI26-1KT, Merck, USA) were seeded in a 24-well plate and allowed to adhere. rhIL-2/rhIL-15 NK cells were co-cultured with ARPE-19 at 3:1, 5:1 (effector to target ratio) at 37°C for 24h. PKH26 stained cells were then stained using Annexin V-FITC/7-AAD Apoptosis Kit (E-CK-A212, Elabscience, USA) for 15 minutes according to manufacturer’s protocol.

### Live cell imaging using Nanolive^TM^ (3D cell Explorer)

ARPE-19 cells (1X10^5^ cells) were seeded on glass bottom (35 mm, CellVis μ-Dish) and incubated at 37°C/5% CO_2._ ARPE-19 cells were co-incubated with activated NK cells stained with PKH26 red cell linker membrane labelling dye (MIDI26-1KT, Merck, USA) at 9:1 (NK cell:ARPE-19 ratio). Live-cell imaging was performed using Nanolive^TM^ 3D Cell Explorer and acquisition by Steve software v1.8.2, Nanolive^TM^ in a temperature-controlled incubation chamber for 16 hours immediately after addition of activated NK cells. Holotomography will allow us to observe changes in membrane, cytosol and nuclear organization and cell-cell interactions at 60X magnification^81^. Images were obtained at 5 minutes intervals. Post-acquisition, 2D tiffs and AVI exports were done using Steve software. AVI exports were done at 10 frames per second (fps).

### Gene expression by Real time Quantitative PCR

Total RNA was isolated from iRPE cells (with and without treatments) using TRIzol according to manufacturer’s instructions (Invitrogen, USA). The concentration of RNA was quantified using BioPhotometer plus (Eppendorf, USA) and cDNA synthesized using iScript kit (Bio-Rad, Philadelphia, USA). The expression of target gene like IL-1β was measured on CFX connect real-time PCR detection system (Bio-Rad, Philadelphia, PA, USA) and normalised to GAPDH (housekeeping gene).

### Western Blotting

The whole cell lysate for immunoblotting was prepared as previously described^82^ Briefly, cells were lysed on ice with RIPA buffer by intermittent vortexing and clarified lysate collected by centrifugation at 13000rpm for 15minutes/4°C. Protein concentration was measured using Bicinchoninic Acid (BCA) Protein Assay (G Biosciences, USA). Whole cell lysates (20μg) were resolved on 12% sodium dodecyl sulfate polyacrylamide gel electrophoresis (SDS-PAGE) gel. The separated proteins were then transferred onto a PVDF membrane and blot was blocked in 5% non-fat milk prepared in 0.1% TRIS-buffered saline-Tween-20 at 4°C overnight. Blots were then incubated with anti-Gasdermin E and anti-GAPDH (Housekeeping protein) for 2h at room temperature. The membranes were washed, probed with anti-rabbit conjugated to HRP for 2h at room temperature. Clarity ECL Western blotting substrate (Bio-Rad) was added to washed membranes and resulting chemiluminescence was measured using Image quant (Amersham Image Quant 800, Cytiva).

### Retinal histology and immunofluorescence of donor eye

Human donor eye was fixed in Neutral buffer formalin for 48h followed by paraffin embedding. 4um thick retinal sections were prepared and used for H&E staining. The stained retinal sections were imaged with Olympus CKX53. Subsequently, for immunostaining, 4um thick sections were deparaffinized and antigen retrieval performed using citrate buffer pH 6. The sections were blocked with 1% BSA /0.1% Triton X-100 for 1h at room temperature, stained overnight at 4C with anti-Granzyme B antibody (Mouse Antibody, 1:100 dilution, 674602, Biolegend, USA) and anti-Occludin antibody (Rabbit Antibody, 1:100 dilution, #91131, Cell Signaling Technology) and incubated with corresponding secondary antibodies (1:2000 dilution) for 1h at room temperature. Following washing with 1xPBST (2 times; 5 mins each), the retinal sections were mounted with DAPI fluoroshield for imaging on Olympus CKX53.

### Intravitreal injection of adenosine

10 month old *Cryba1* cKO mouse were intravitreally injected (PMID: 31552301) with adenosine (Sigma Aldrich, A4036-5G) at a dose of 4μM, once for three consecutive weeks, in one eye whereas the contralateral eyes was injected with PBS (carrier). After one month from the first injection, mice were euthanized, aqueous humor (immunophenotyping and ELISA) and RPE (western blot) were collected for experiments.

### Statistical analysis

Shapiro-Wilk normality test followed by Mann-Whitney test, Kruskal-Wallis test followed by Dunn multiple comparison test, Ordinary one way ANOVA followed by Tukey’s multiple comparison test and Spearman Rank Correlation analysis was performed to determine the distribution of the data, parametric/non-parametric comparison between groups and association between immune cells and effector factors, respectively. Statistical analyses were performed with GraphPad Prism 10.1 (GraphPad Software, Inc., La Jolla, CA, USA), Orange and MedCalc® Version 12.5 (MedCalc Software, Ostend, Belgium). p value< 0.05 was considered statistically significant.

## ACKNOWLEDGEMENTS

Narayana Netralaya Foundation, Bangalore, India, provided the financial support for the study. The authors acknowledge the assistance of clinical fellows Prashanti C V, Satyanarayanamurthy Ayyori, Aishwariya Joshi who assisted with the clinical classification of patient aqueous humor. We thank Packiyaraj Pandian, Shubham Pawar, and Steffi Varghese for their help with processing human aqueous humor samples and donor eyeballs; Sourab Shetty for his help with the paraffin block preparation of donor eyeballs; Ravikiran and Pradeep K for their help with processing of donor retina for untargeted metabolomics. We would like to thank J. Samuel Zigler, Jr. for his help during the preparation of this manuscript. This work was also supported by 2019 Roche Collaborative Research Fellowship on behalf of ARVO Foundation for Eye Research (SS), NIH 1R01EY031594-01A1 (DS & JTH), NIH 1R01EY032516-01A1 (DS), The Johns Hopkins University School of Medicine start-up funds (DS), Edward N. & Della L. Thome Memorial Foundation Awards Program in Age-Related Macular Degeneration (DS), BrightFocus Foundation Postdoctoral Fellowship on Macular Degeneration (to SG), NIH K99/R00 (1K99EY033421-01A1) Pathway to Independence award from the National Eye Institute, NIH (to SG), P30 core award (EY001765) from the National Eye Institute, NIH to the Wilmer Eye Institute, the Johns Hopkins University School of Medicine, and unrestricted funds from The Research to Prevent Blindness Inc., NY to the Wilmer Eye Institute, the Johns Hopkins University School of Medicine. DS is the Frieda Derdeyn Bambas Professor of Ophthalmology.

## AUTHOR CONTRIBUTIONS

Conceptualization and project administration: Arkasubhra Ghosh, Debasish Sinha

Study design and Methodology: Arkasubhra Ghosh, Debasish Sinha, Swaminathan Sethu, J. Jiang Qian, Vrushali Deshpande, Koushik Chakrabarty.

Conducted experiments and analysis-Archana Padmanabhan Nair, Sayan Ghosh, Ying Xin, Vishnu Suresh Babu, Machiraju Praveen, Ganesh Ram Sahu, Tanuja Arun Vaidya, Jayasree Debnath, Karthik Raja.

Funding acquisition: Arkasubhra Ghosh, Debasish Sinha, Swaminathan Sethu

Resources and clinical data curation: Santhosh Gopikrishna Gadde, Thirumalesh M B, Naren Shetty, Aishwarya Saxena, Rohit Shetty.

Wrote the manuscript: Archana Padmanabhan Nair, Sayan Ghosh, Arkasubhra Ghosh, Debasish Sinha.

Reviewed & edited the manuscript: Swaminathan Sethu, Stacey Hose, James T. Handa, Arkasubhra Ghosh, Debasish Sinha.

## REFERENCES

1 Flaxman, S. R. et al. Global causes of blindness and distance vision impairment 1990-2020: a systematic review and meta-analysis. Lancet Glob Health 5, e1221–e1234 (2017). 10.1016/S2214-109X(17)30393-5

2 Somasundaran, S., Constable, I. J., Mellough, C. B. & Carvalho, L. S. Retinal pigment epithelium and age-related macular degeneration: A review of major disease mechanisms. Clin Exp Ophthalmol 48, 1043–1056 (2020). 10.1111/ceo.13834

3 Boopathiraj, N., Wagner, I. V., Dorairaj, S. K., Miller, D. D. & Stewart, M. W. Recent Updates on the Diagnosis and Management of Age-Related Macular Degeneration. Mayo Clin Proc Innov Qual Outcomes 8, 364–374 (2024). 10.1016/j.mayocpiqo.2024.05.003

4 Nebbioso, M. et al. Therapeutic Approaches with Intravitreal Injections in Geographic Atrophy Secondary to Age-Related Macular Degeneration: Current Drugs and Potential Molecules. Int J Mol Sci 20 (2019). 10.3390/ijms20071693

5 Davis, M. D. et al. The Age-Related Eye Disease Study severity scale for age-related macular degeneration: AREDS Report No. 17. Arch Ophthalmol 123, 1484–1498 (2005). 10.1001/archopht.123.11.1484

6 Kerur, N. et al. cGAS drives noncanonical-inflammasome activation in age-related macular degeneration. Nat Med 24, 50–61 (2018). 10.1038/nm.4450

7 Wang, M., Su, S., Jiang, S., Sun, X. & Wang, J. Role of amyloid beta-peptide in the pathogenesis of age-related macular degeneration. BMJ Open Ophthalmol 6, e000774 (2021). 10.1136/bmjophth-2021-000774

8 Wang, S. et al. Autophagy Dysfunction, Cellular Senescence, and Abnormal Immune-Inflammatory Responses in AMD: From Mechanisms to Therapeutic Potential. Oxid Med Cell Longev 2019, 3632169 (2019). 10.1155/2019/3632169

9 Gupta, U. et al. Increased LCN2 (lipocalin 2) in the RPE decreases autophagy and activates inflammasome-ferroptosis processes in a mouse model of dry AMD. Autophagy 19, 92–111 (2023). 10.1080/15548627.2022.2062887

10 Lambert, N. G. et al. Risk factors and biomarkers of age-related macular degeneration. Prog Retin Eye Res 54, 64–102 (2016). 10.1016/j.preteyeres.2016.04.003

11 Ambati, J. & Fowler, B. J. Mechanisms of age-related macular degeneration. Neuron 75, 26–39 (2012). 10.1016/j.neuron.2012.06.018

12 Bowes Rickman, C., Farsiu, S., Toth, C. A. & Klingeborn, M. Dry age-related macular degeneration: mechanisms, therapeutic targets, and imaging. Invest Ophthalmol Vis Sci 54, ORSF68–80 (2013). 10.1167/iovs.13-12757

13 Knickelbein, J. E., Chan, C. C., Sen, H. N., Ferris, F. L. & Nussenblatt, R. B. Inflammatory Mechanisms of Age-related Macular Degeneration. Int Ophthalmol Clin 55, 63–78 (2015). 10.1097/IIO.0000000000000073

14 Copland, D. A., Theodoropoulou, S., Liu, J. & Dick, A. D. A Perspective of AMD Through the Eyes of Immunology. Invest Ophthalmol Vis Sci 59, AMD83–AMD92 (2018). 10.1167/iovs.18-23893

15 Allingham, M. J., Loksztejn, A., Cousins, S. W. & Mettu, P. S. Immunological Aspects of Age-Related Macular Degeneration. Adv Exp Med Biol 1256, 143–189 (2021). 10.1007/978-3-030-66014-7_6

16 Charbel Issa, P., Chong, N. V. & Scholl, H. P. The significance of the complement system for the pathogenesis of age-related macular degeneration - current evidence and translation into clinical application. Graefes Arch Clin Exp Ophthalmol 249, 163–174 (2011). 10.1007/s00417-010-1568-6

17 Cruz-Pimentel, M. & Wu, L. Complement Inhibitors for Advanced Dry Age-Related Macular Degeneration (Geographic Atrophy): Some Light at the End of the Tunnel? J Clin Med 12 (2023). 10.3390/jcm12155131

18 Boyce, M. et al. Microglia-Neutrophil Interactions Drive Dry AMD-like Pathology in a Mouse Model. Cells 11 (2022). 10.3390/cells11223535

19 Ghosh, S. et al. Neutrophils homing into the retina trigger pathology in early age-related macular degeneration. Commun Biol 2, 348 (2019). 10.1038/s42003-019-0588-y

20 Roubeix, C. et al. Splenic monocytes drive pathogenic subretinal inflammation in age-related macular degeneration. J Neuroinflammation 21, 22 (2024). 10.1186/s12974-024-03011-z

21 Levy, O. et al. Apolipoprotein E promotes subretinal mononuclear phagocyte survival and chronic inflammation in age-related macular degeneration. EMBO Mol Med 7, 211–226 (2015). 10.15252/emmm.201404524

22 Sennlaub, F. et al. CCR2(+) monocytes infiltrate atrophic lesions in age-related macular disease and mediate photoreceptor degeneration in experimental subretinal inflammation in Cx3cr1 deficient mice. EMBO Mol Med 5, 1775–1793 (2013). 10.1002/emmm.201302692

23 Daftarian, N. et al. Peripheral blood CD163(+) monocytes and soluble CD163 in dry and neovascular age-related macular degeneration. FASEB J 34, 8001–8011 (2020). 10.1096/fj.201901902RR

24 Cruz-Guilloty, F. et al. T cells and macrophages responding to oxidative damage cooperate in pathogenesis of a mouse model of age-related macular degeneration. PLoS One 9, e88201 (2014). 10.1371/journal.pone.0088201

25 Bhutto, I. A. et al. Increased choroidal mast cells and their degranulation in age-related macular degeneration. Br J Ophthalmol 100, 720–726 (2016). 10.1136/bjophthalmol-2015-308290

26 Chang, Y. H. et al. Protective role of IL-17-producing gammadelta T cells in a laser-induced choroidal neovascularization mouse model. J Neuroinflammation 20, 279 (2023). 10.1186/s12974-023-02952-1

27 Qi, C. & Liu, Q. Natural killer cells in aging and age-related diseases. Neurobiol Dis 183, 106156 (2023). 10.1016/j.nbd.2023.106156

28 Xiao, J. et al. High-Fat Diet Alters the Retinal Pigment Epithelium and Choroidal Transcriptome in the Absence of Gut Microbiota. Cells 11 (2022). 10.3390/cells11132076

29 Matsubara, J. A. et al. Retinal Distribution and Extracellular Activity of Granzyme B: A Serine Protease That Degrades Retinal Pigment Epithelial Tight Junctions and Extracellular Matrix Proteins. Front Immunol 11, 574 (2020). 10.3389/fimmu.2020.00574

30 Cousins, S. W., Trattler, W. B. & Streilein, J. W. Immune privilege and suppression of immunogenic inflammation in the anterior chamber of the eye. Curr Eye Res 10, 287–297 (1991). 10.3109/02713689108996334

31 Taylor, A. W. Ocular immunosuppressive microenvironment. Chem Immunol Allergy 92, 71–85 (2007). 10.1159/000099255

32 Niederkorn, J. Y. Mechanisms of immune privilege in the eye and hair follicle. J Investig Dermatol Symp Proc 8, 168–172 (2003). 10.1046/j.1087-0024.2003.00803.x

33 Keino, H., Horie, S. & Sugita, S. Immune Privilege and Eye-Derived T-Regulatory Cells. J Immunol Res 2018, 1679197 (2018). 10.1155/2018/1679197

34 Taylor, A. W. & Ng, T. F. Negative regulators that mediate ocular immune privilege. J Leukoc Biol (2018). 10.1002/JLB.3MIR0817-337R

35 Du, Y. & Yan, B. Ocular immune privilege and retinal pigment epithelial cells. J Leukoc Biol 113, 288–304 (2023). 10.1093/jleuko/qiac016

36 Wong, J. H. C. et al. Exploring the pathogenesis of age-related macular degeneration: A review of the interplay between retinal pigment epithelium dysfunction and the innate immune system. Front Neurosci 16, 1009599 (2022). 10.3389/fnins.2022.1009599

37 Spindler, J., Zandi, S., Pfister, I. B., Gerhardt, C. & Garweg, J. G. Cytokine profiles in the aqueous humor and serum of patients with dry and treated wet age-related macular degeneration. PLoS One 13, e0203337 (2018). 10.1371/journal.pone.0203337

38 Collin, J. et al. Single-cell RNA sequencing reveals transcriptional changes of human choroidal and retinal pigment epithelium cells during fetal development, in healthy adult and intermediate age-related macular degeneration. Hum Mol Genet 32, 1698–1710 (2023). 10.1093/hmg/ddad007

39 Ambati, J., Atkinson, J. P. & Gelfand, B. D. Immunology of age-related macular degeneration. Nat Rev Immunol 13, 438–451 (2013). 10.1038/nri3459

40 Shang, P. et al. The amino acid transporter SLC36A4 regulates the amino acid pool in retinal pigmented epithelial cells and mediates the mechanistic target of rapamycin, complex 1 signaling. Aging Cell 16, 349–359 (2017). 10.1111/acel.12561

41 Ghosh, S. et al. A Role for betaA3/A1-Crystallin in Type 2 EMT of RPE Cells Occurring in Dry Age-Related Macular Degeneration. Invest Ophthalmol Vis Sci 59, AMD104–AMD113 (2018). 10.1167/iovs.18-24132

42 Bizymi, N., Matthaiou, A. M., Mavroudi, I., Batsali, A. & Papadaki, H. A. Immunomodulatory actions of myeloid-derived suppressor cells in the context of innate immunity. Innate Immun 30, 2–10 (2024). 10.1177/17534259231215581

43 Kamao, H. et al. Characterization of human induced pluripotent stem cell-derived retinal pigment epithelium cell sheets aiming for clinical application. Stem Cell Reports 2, 205–218 (2014). 10.1016/j.stemcr.2013.12.007

44 Ablonczy, Z. et al. Human retinal pigment epithelium cells as functional models for the RPE in vivo. Invest Ophthalmol Vis Sci 52, 8614–8620 (2011). 10.1167/iovs.11-8021

45 Dunn, K. C., Aotaki-Keen, A. E., Putkey, F. R. & Hjelmeland, L. M. ARPE-19, a human retinal pigment epithelial cell line with differentiated properties. Exp Eye Res 62, 155–169 (1996). 10.1006/exer.1996.0020

46 Hazim, R. A., Volland, S., Yen, A., Burgess, B. L. & Williams, D. S. Rapid differentiation of the human RPE cell line, ARPE-19, induced by nicotinamide. Exp Eye Res 179, 18–24 (2019). 10.1016/j.exer.2018.10.009

47 Hu, Y. et al. The multifaceted roles of GSDME-mediated pyroptosis in cancer: therapeutic strategies and persisting obstacles. Cell Death Dis 14, 836 (2023). 10.1038/s41419-023-06382-y

48 Kumar, V. Adenosine as an endogenous immunoregulator in cancer pathogenesis: where to go? Purinergic Signal 9, 145–165 (2013). 10.1007/s11302-012-9349-9

49 Ohta, A. A Metabolic Immune Checkpoint: Adenosine in Tumor Microenvironment. Front Immunol 7, 109 (2016). 10.3389/fimmu.2016.00109

50 Pasquini, S., Contri, C., Borea, P. A., Vincenzi, F. & Varani, K. Adenosine and Inflammation: Here, There and Everywhere. Int J Mol Sci 22 (2021). 10.3390/ijms22147685

51 Hoskin, D. W., Mader, J. S., Furlong, S. J., Conrad, D. M. & Blay, J. Inhibition of T cell and natural killer cell function by adenosine and its contribution to immune evasion by tumor cells (Review). Int J Oncol 32, 527–535 (2008).

52 Chambers, A. M. et al. Adenosinergic Signaling Alters Natural Killer Cell Functional Responses. Front Immunol 9, 2533 (2018). 10.3389/fimmu.2018.02533

53 Chambers, A. M. & Matosevic, S. Immunometabolic Dysfunction of Natural Killer Cells Mediated by the Hypoxia-CD73 Axis in Solid Tumors. Front Mol Biosci 6, 60 (2019). 10.3389/fmolb.2019.00060

54 Chen, M. & Xu, H. Parainflammation, chronic inflammation, and age-related macular degeneration. J Leukoc Biol 98, 713–725 (2015). 10.1189/jlb.3RI0615-239R

55 Mullins, R. F., Russell, S. R., Anderson, D. H. & Hageman, G. S. Drusen associated with aging and age-related macular degeneration contain proteins common to extracellular deposits associated with atherosclerosis, elastosis, amyloidosis, and dense deposit disease. FASEB J 14, 835–846 (2000).

56 Wang, J. et al. ATAC-Seq analysis reveals a widespread decrease of chromatin accessibility in age-related macular degeneration. Nat Commun 9, 1364 (2018). 10.1038/s41467-018-03856-y

57 Borchert, G. A. et al. The Role of Inflammation in Age-Related Macular Degeneration-Therapeutic Landscapes in Geographic Atrophy. Cells 12 (2023). 10.3390/cells12162092

58 Kauppinen, A., Paterno, J. J., Blasiak, J., Salminen, A. & Kaarniranta, K. Inflammation and its role in age-related macular degeneration. Cell Mol Life Sci 73, 1765–1786 (2016). 10.1007/s00018-016-2147-8

59 Rosa, J. G. S., Disner, G. R., Pinto, F. J., Lima, C. & Lopes-Ferreira, M. Revisiting Retinal Degeneration Hallmarks: Insights from Molecular Markers and Therapy Perspectives. Int J Mol Sci 24 (2023). 10.3390/ijms241713079

60 Chen, S., Zhu, H. & Jounaidi, Y. Comprehensive snapshots of natural killer cells functions, signaling, molecular mechanisms and clinical utilization. Signal Transduct Target Ther 9, 302 (2024). 10.1038/s41392-024-02005-w

61 Naujoks, W. et al. Characterization of Surface Receptor Expression and Cytotoxicity of Human NK Cells and NK Cell Subsets in Overweight and Obese Humans. Front Immunol 11, 573200 (2020). 10.3389/fimmu.2020.573200

62 Wang, H. Y. et al. Dynamic changes of phenotype and function of natural killer cells in peripheral blood before and after thermal ablation of hepatitis B associated hepatocellular carcinoma and their correlation with tumor recurrence. BMC Cancer 23, 486 (2023). 10.1186/s12885-023-10823-4

63 Lee, G. et al. NK cells from COVID-19 positive patients exhibit enhanced cytotoxic activity upon NKG2A and KIR2DL1 blockade. Front Immunol 14, 1022890 (2023). 10.3389/fimmu.2023.1022890

64 Luo, Y. et al. Single-cell RNA Sequencing Identifies Natural Kill Cell-Related Transcription Factors Associated With Age-Related Macular Degeneration. Evol Bioinform Online 20, 11769343241272413 (2024). 10.1177/11769343241272413

65 Dong, X. et al. Natural killer cells promote neutrophil extracellular traps and restrain macular degeneration in mice. Sci Transl Med 16, eadi6626 (2024). 10.1126/scitranslmed.adi6626

66 Walsh, M. J. et al. IFNgamma is a central node of cancer immune equilibrium. Cell Rep 42, 112219 (2023). 10.1016/j.celrep.2023.112219

67 Holbrook, J. et al. Natural killer cells have an activated profile in early Parkinson’s disease. J Neuroimmunol 382, 578154 (2023). 10.1016/j.jneuroim.2023.578154

68 Antonioli, L., Pacher, P. & Hasko, G. Adenosine and inflammation: it’s time to (re)solve the problem. Trends Pharmacol Sci 43, 43–55 (2022). 10.1016/j.tips.2021.10.010

69 Losenkova, K. et al. CD73 controls ocular adenosine levels and protects retina from light-induced phototoxicity. Cell Mol Life Sci 79, 152 (2022). 10.1007/s00018-022-04187-4

70 Ye, S. S., Tang, Y. & Song, J. T. ATP and Adenosine in the Retina and Retinal Diseases. Front Pharmacol 12, 654445 (2021). 10.3389/fphar.2021.654445

71 Bruno, A., Mortara, L., Baci, D., Noonan, D. M. & Albini, A. Myeloid Derived Suppressor Cells Interactions With Natural Killer Cells and Pro-angiogenic Activities: Roles in Tumor Progression. Front Immunol 10, 771 (2019). 10.3389/fimmu.2019.00771

72 Wang, L. et al. Mechanisms of PANoptosis and relevant small-molecule compounds for fighting diseases. Cell Death Dis 14, 851 (2023). 10.1038/s41419-023-06370-2

73 Du, T. et al. Pyroptosis, metabolism, and tumor immune microenvironment. Clin Transl Med 11, e492 (2021). 10.1002/ctm2.492

74 Yang, X. et al. IFN-gamma Facilitates Corneal Epithelial Cell Pyroptosis Through the JAK2/STAT1 Pathway in Dry Eye. Invest Ophthalmol Vis Sci 64, 34 (2023). 10.1167/iovs.64.3.34

75 Shang, P. et al. betaA3/A1-crystallin regulates apical polarity and EGFR endocytosis in retinal pigmented epithelial cells. Commun Biol 4, 850 (2021). 10.1038/s42003-021-02386-6

76 Valapala, M. et al. Lysosomal-mediated waste clearance in retinal pigment epithelial cells is regulated by CRYBA1/betaA3/A1-crystallin via V-ATPase-MTORC1 signaling. Autophagy 10, 480–496 (2014). 10.4161/auto.27292

77 Yang, Y. et al. New sample preparation approach for mass spectrometry-based profiling of plasma results in improved coverage of metabolome. J Chromatogr A 1300, 217–226 (2013). 10.1016/j.chroma.2013.04.030

78 Xin, Y. et al. LRLoop: a method to predict feedback loops in cell-cell communication. Bioinformatics 38, 4117–4126 (2022). 10.1093/bioinformatics/btac447

79 Cabello-Aguilar, S. et al. SingleCellSignalR: inference of intercellular networks from single-cell transcriptomics. Nucleic Acids Res 48, e55 (2020). 10.1093/nar/gkaa183

80 Foltz, L. P. & Clegg, D. O. Rapid, Directed Differentiation of Retinal Pigment Epithelial Cells from Human Embryonic or Induced Pluripotent Stem Cells. J Vis Exp (2017). 10.3791/56274

81 Nadjar, J. et al. An optogenetic approach to control and monitor inflammasome activation. bioRxiv, 2023.2007.2025.550490 (2023). 10.1101/2023.07.25.550490

82 Panigrahi, T. et al. Genistein-Calcitriol Mitigates Hyperosmotic Stress-Induced TonEBP, CFTR Dysfunction, VDR Degradation and Inflammation in Dry Eye Disease. Clin Transl Sci 14, 288–298 (2021). 10.1111/cts.12858

