## Supplementary Figures, Legends and Methods for "Attenuated adenosine mediated immune-dampening increases natural killer cell activity in early age-related macular degeneration"

### Supplemental Figures

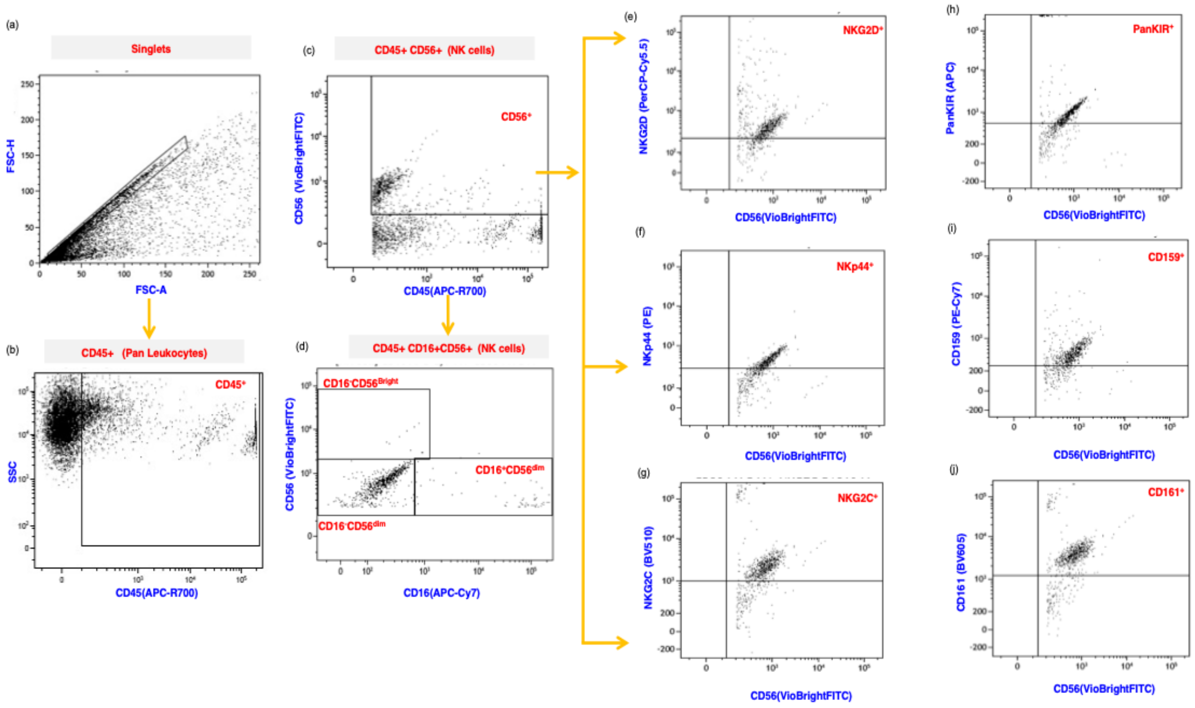


**Supplemental Figure 1 I** Gating strategy to identify NK cell subsets in early AMD subjects AH and human donor retina. Representative images showing gating strategy used to determine the NK cell subsets. The outlined region in (a) and (b) represents singlets and leukocyte population respectively. (c) The outlined region shows cells double positive for CD56 (VioBright-FITC) and CD45 (APC-R700). (d) This plot with CD16 (APC-Cy7) and CD56 (VioBright-FITC) shows CD56+ Natural killer cells sub-divided based on expression of CD16 and CD56 into CD16-CD56$^{bright}$ and CD16+CD56$^{dim}$ representing cytokine producing and cytotoxic forms of NK populations, respectively. An intermediate population of CD16-CD56+ cells was shown. (e,f,g) These panels show CD56 (VioBright-FITC) versus NKG2D (PerCP-Cy5.5), NKp44 (PE) and NKG2C (BV510) where cells marked double positive for respective markers are NK cell subsets expressing activating receptors. (h,i,j) These panels show CD56 (VioBright-FITC) versus PanKIR2D (APC), CD159a (PE-Cy7) and CD161 (BV605) where cells double positive are NK subsets expressing inhibitory receptors. The percentages of immune cells were computed using the number of positive cells for each marker to total number of cells acquired in each sample, while the percentage of NK cells expressing activating and inhibitory were computed using number of positive cells for each marker to total NK cell number obtained per sample .

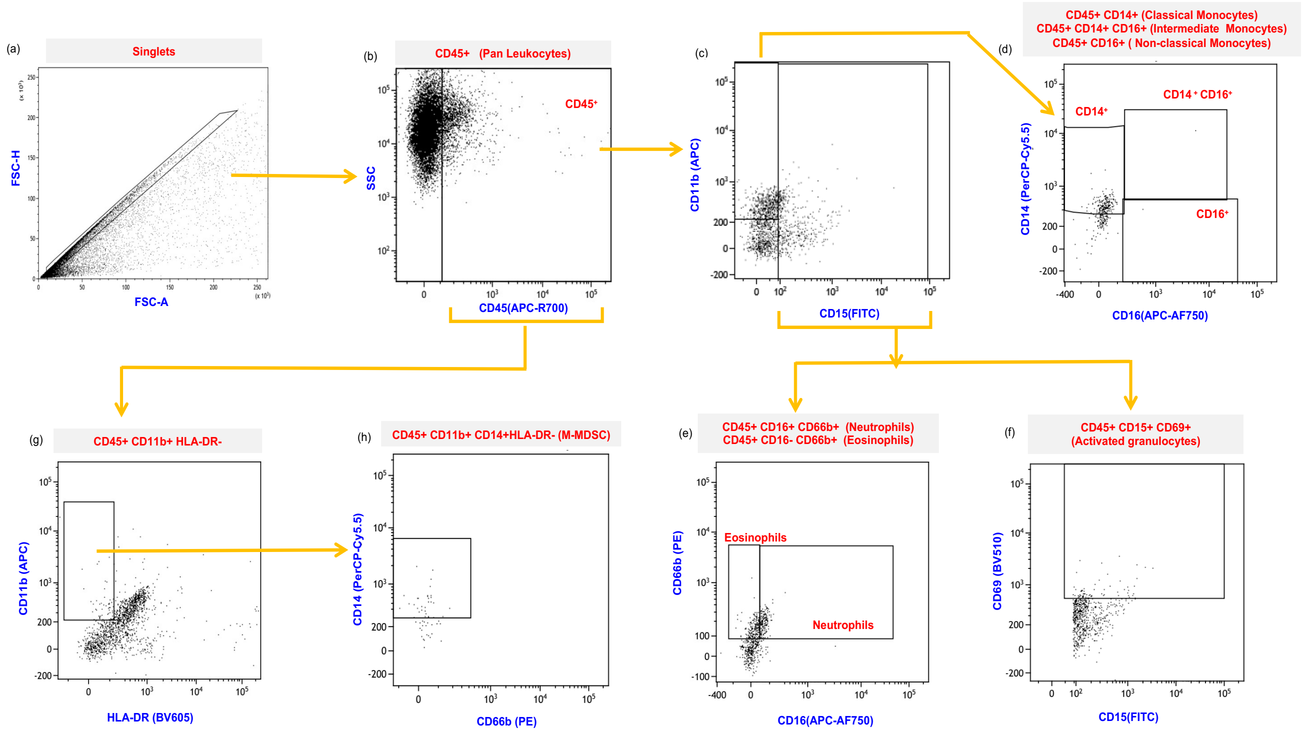


**Supplemental Figure 2 I** Gating strategy to identify Myeloid cell subsets in AH of early AMD subjects. Representative images show gating strategy used to determine the myeloid cell subsets. (a) The outlined region in FSC-H versus FSC-A plot represents Singlets. (b) The marked region CD45 (APC-R700) versus Side Scatter (SSC) represents total leukocyte population. (c) The outlined region in CD15 (FITC) versus CD11b (APC) plot represents Granulocytes (right region) and Non-granulocytes (left region).(d)The plot with CD14 (PerCP-Cy5.5) versus CD16 (APC-AF750), represents classical monocytes(CD14+), intermediate monocytes (CD14+CD16+) and non-classical monocytes (CD16+), within non-granulocyte population.(e) Within granulocyte population, the right region in the CD16 (APC-AF750) versus CD66b (PE) plot shows cells double positive for CD16 and CD66b, representing Neutrophils, while left region shows cells positive for CD66b alone, representing Eosinophils. (f) The upper region in this panel shows cells positive for both CD15 (FITC) and CD69 (BV510) representing Activated granulocytes. (g) This panel shows CD11b (APC) versus HLA-DR(BV605), where cells negative for HLA-DR and positive for CD11b were gated in panel h. (h) This panel shows CD14 (PerCP-Cy5.5) versus CD66b (PE), where cells positive for CD14 (PerCP-Cy5.5) represent monocytic-Myeloid derived suppressor (M-MDSC) cells. The percentages of immune cells were computed using number of positive cells for each marker to total number of cells acquired in each sample.

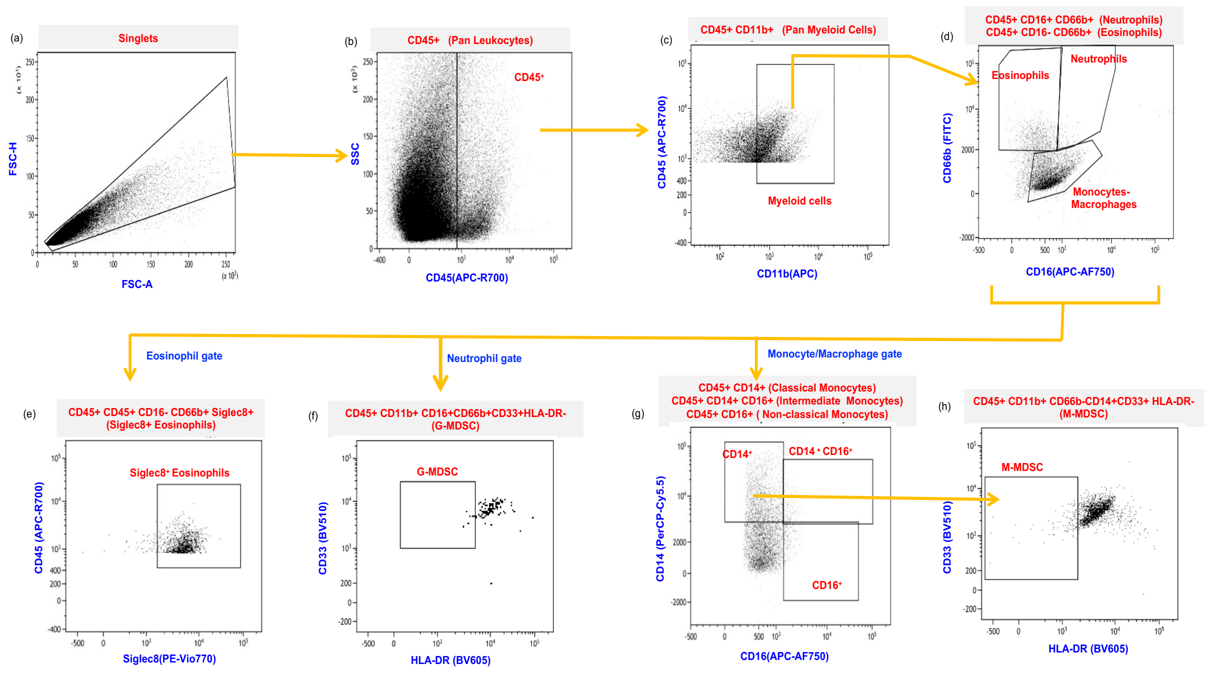


**Supplemental Figure 3 I** Gating strategy to identify Myeloid cell subsets in human donor retinae. Representative images show gating strategy used to determine the myeloid cell subsets. (a) The outlined region in FSC-H versus FSC-A plot represents singlet population. (b) The marked region CD45 (APC-R700) versus Side Scatter (SSC) represents total leukocyte population. (c) The outlined region in CD45 (APC-R700) versus CD11b (APC) plot represents total myeloid cells. (d)The plot with CD66b (FITC) versus CD16 (APC-AF750), represents neutrophils, cells double positive for CD16 and CD66b which is right region in the plot; while left region shows cells positive for CD66b without CD16 expression, representing eosinophils. The region in the same plot with cells positive for CD16 and no expression is seen for CD66b, represents monocytes/macrophages. (e) The plot with CD45 (APC-R700) versus Siglec 8 (PE-Vio770), represents Siglec 8+ eosinophils (within the eosinophil population). (f) This panel shows CD33 (BV510) versus HLA-DR (BV605) within neutrophil population, where cells expressing CD33 and negative for HLA-DR represent granulocytic-Myeloid derived suppressor (G-MDSC) cells. (g) The outlined regions in CD14 (PerCP-Cy5.5) versus CD16 (APC-AF750) panel represents classical monocytes(CD14+), intermediate monocytes (CD14+CD16+) and non-classical monocytes (CD16+), within monocyte/macrophage population.(h) This panel shows CD33 (BV510) versus HLA-DR(BV605), where cells negative for HLA-DR and positive for CD33 were gated based on CD14+ monocyte population represents monocytic-Myeloid derived suppressor (M-MDSC) cells. The percentages of immune cells were computed using number of positive cells for each marker to total number of cells acquired in each sample.

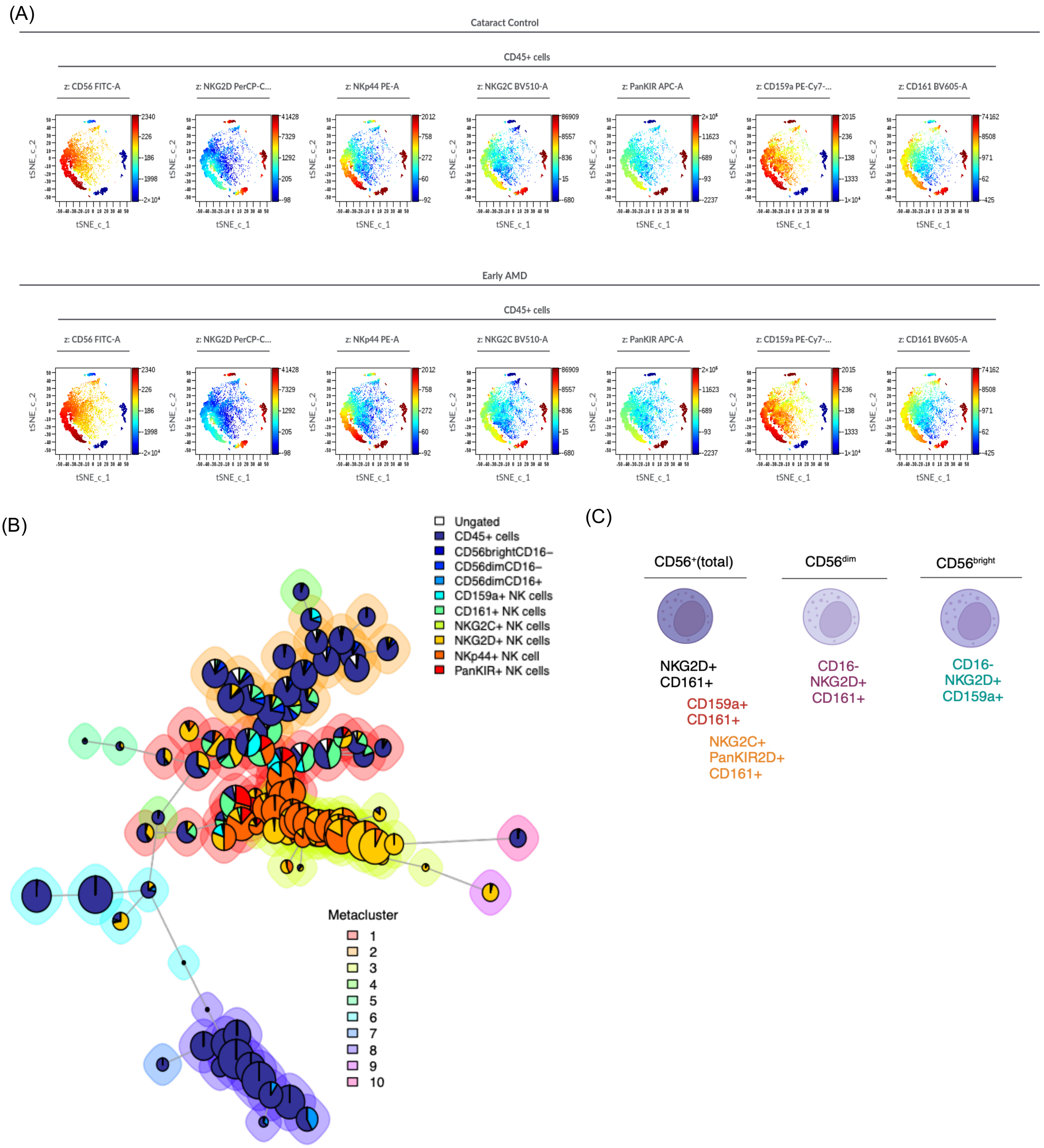
(A)
Cataract Control
CD45+ cells
z: CD56 FITC-A
z: NKG2D PerCP-C...
z: NKp44 PE-A
z: NKG2C BV510-A
z: PanKIR APC-A
z: CD159a PE-Cy7-...
z: CD161 BV605-A
tSNE_c_1
tSNE_c_2
Early AMD
CD45+ cells
z: CD56 FITC-A
z: NKG2D PerCP-C...
z: NKp44 PE-A
z: NKG2C BV510-A
z: PanKIR APC-A
z: CD159a PE-Cy7-...
z: CD161 BV605-A
tSNE_c_1
tSNE_c_2
(B)
Ungated
CD45+ cells
CD56brightCD16–
CD56dimCD16–
CD56dimCD16+
CD159a+ NK cells
CD161+ NK cells
NKG2C+ NK cells
NKG2D+ NK cells
NKp44+ NK cell
PanKIR+ NK cells
Metacluster
1
2
3
4
5
6
7
8
9
10
(C)
CD56+(total)
CD56dim
CD56bright
NKG2D+
CD161+
CD159a+
CD161+
NKG2C+
PanKIR2D+
CD161+
CD16-
NKG2D+
CD161+
CD16-
NKG2D+
CD159a+

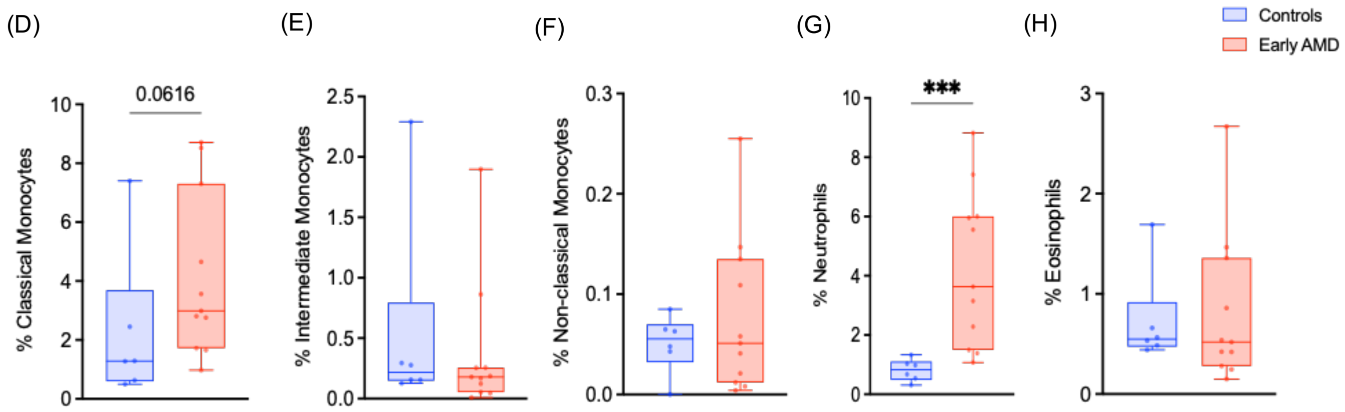


**Supplemental Figure 4 I** NK cell clustering and Myeloid cell proportions identified in early AMD aqueous humor of pooled subjects. (A) tSNE-CUDA analysis of NK cell subsets (activating and inhibitory receptors) in Cytobank. (B) FLOWSOM based clustering in Cytobank identified 10 metaclusters (cluster 1,2,6,8 focused on NK cells) (C) Illustration summarizing diverse NK cell subsets in human AH with phenotypes including CD56+NKG2D+CD161+, CD56+CD161+CD159+, CD56$^{+}$NKG2C$^{+}$PanKIR2D$^{+}$CD161$^{+}$, CD56$^{dim}$CD16$^{-}$NKG2D$^{+}$CD161$^{+}$ and CD56$^{bright}$CD16$^{-}$NKG2D$^{+}$CD159a$^{+}$. Box and whisker plots show a trend with CD45+CD14+ (classical monocytes) and neutrophils (D,G) and no changes in CD45+CD14+CD16+ (Intermediate monocytes), CD45+CD14-CD16+ (Non-Classical Monocytes) and CD45+CD16-CD66b+ (Eosinophils) (E,F,H) in pooled aqueous humor of cataract control and early AMD subjects. Graphs show Min to Max (all points); ***$P < 0.001$, Mann–Whitney test.

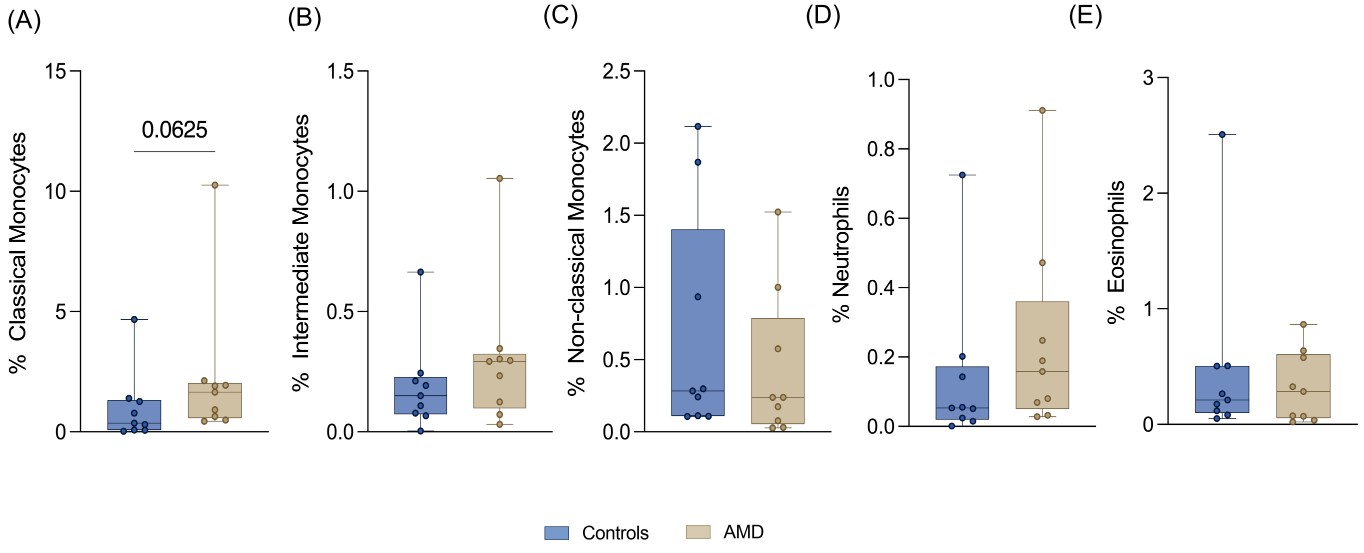


**Supplemental Figure 5 I** Myeloid cell proportions in control and AMD donor retinae. (A-E) Box and whisker plot shows increasing trend in CD45+CD14+ (classical monocytes); no changes seen in CD45+CD14+CD16+ (Intermediate monocytes), CD45+CD14-CD16+ (Non-Classical Monocytes) CD45+CD16+CD66b+ (Neutrophils), or CD45+CD16-CD66b+ (Eosinophils) in AMD donor eye compared to controls. Graphs show Min to Max (all points); Mann–Whitney test.

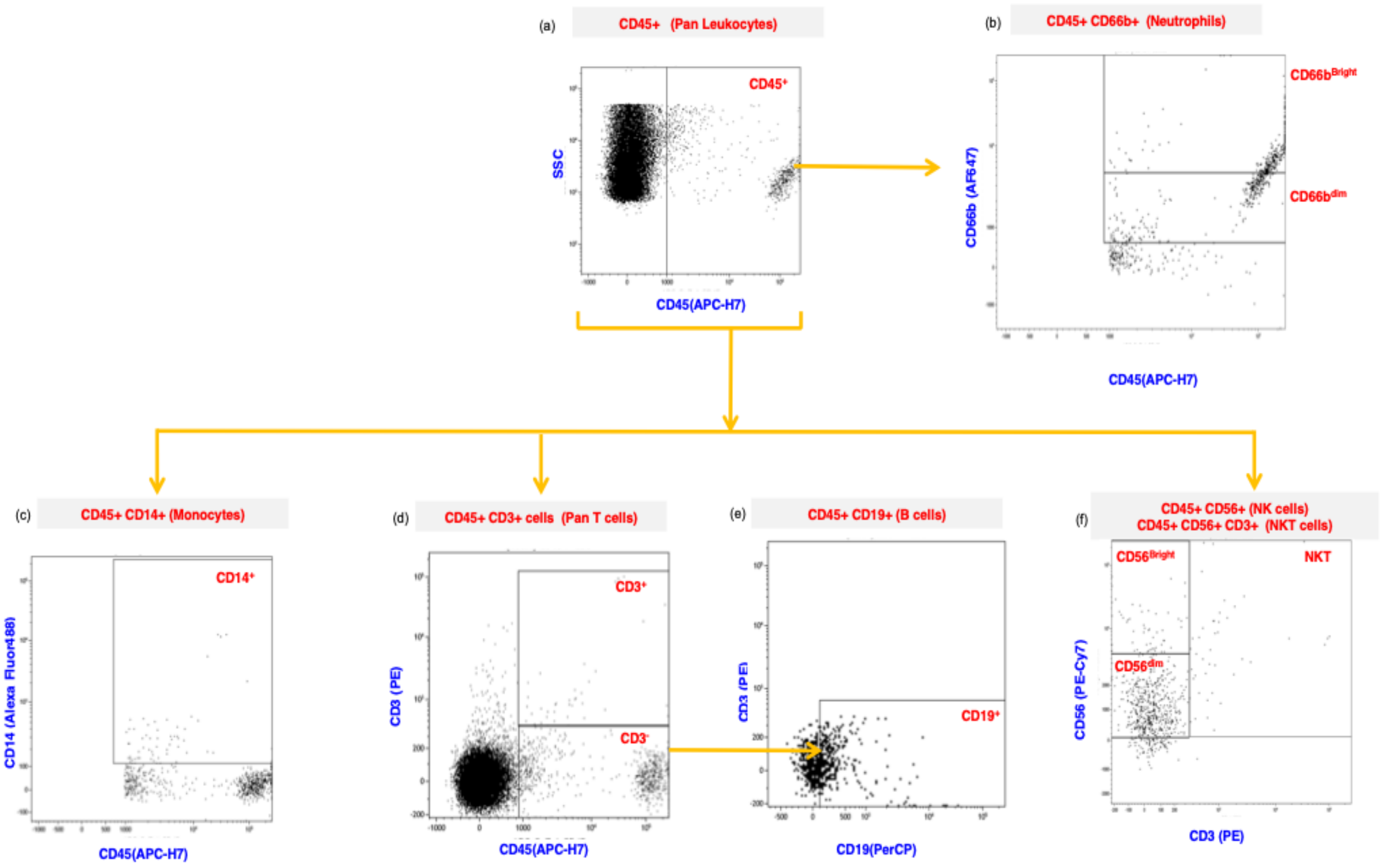


**Supplemental Figure 6 I** Gating strategy for immune cell typing in AMD aqueous humor. Representative images show gating strategy used to determine the immune cell populations in control and early AMD AH samples. (a) The marked region in the CD45 (APC-H7) versus Side Scatter (SSC) plot? represents the leukocyte population. (b) The top right quadrant shows cells double positive for CD66b (AF647) and CD45 (APC-H7). This quadrant was sub-divided based on expression of CD66b into CD66b$^{bright}$ and CD66b$^{dim}$ representing the activated and quiescent neutrophil populations, respectively. (c) The upper right quadrant shows monocytes stained positive for CD14 (Alexa Fluor 488) and CD45 (APC-H7). (d) The upper marked region in this panel shows cells positive for both CD45 (APC-H7) and CD3 (PE) representative of T cells, while the lower region shows cells positive for CD45(APC-H7) and negative for CD3 (PE). (e) This panel shows CD19 (PerCP) versus CD3(PE), where cells negative for CD3 (PE) and positive for CD19 (PerCP) represent B cells. (f) This panel shows CD56 (PE-Cy7) versus CD3(PE), where cells positive for CD56 (PE-Cy7) and negative for CD3 (PE) represent Natural Killer (NK) cells present among the CD45+ cells. The CD56 region is further subdivided into CD56$^{Bright}$ and CD56$^{dim}$ representing cytokine producing and cytotoxic forms of NK populations, respectively. Further, the upper right panel represents cells that are positively stained for both CD3 (PE) and CD56 (PE-Cy7) representing Natural Killer T (NKT) cells. The percentage of each immune cell type was computed using number of positive cells for each marker to total number of cells acquired in each sample.

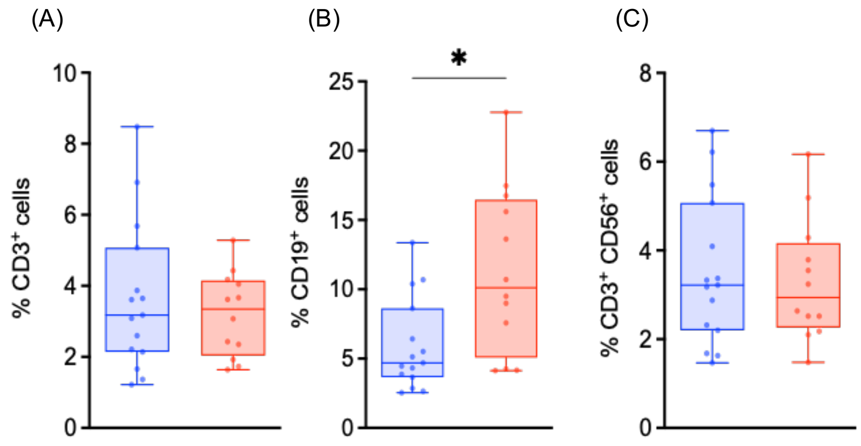


**Supplemental Figure 7 I** Lymphoid cell proportions identified in early AMD aqueous humor (individual subjects). (A,B,C). Box whisker plots show higher proportions of CD45+CD19+ (B cells); no change in CD45+CD3+ (Pan T cells); and CD45+CD3+CD56+ (Natural Killer T – NKT cells) was seen within total cells acquired from aqueous humor of cataract control (n=15) and early AMD (n=12) subjects. Graphs show Min to Max (all points); *P < 0.05, Mann–Whitney test.

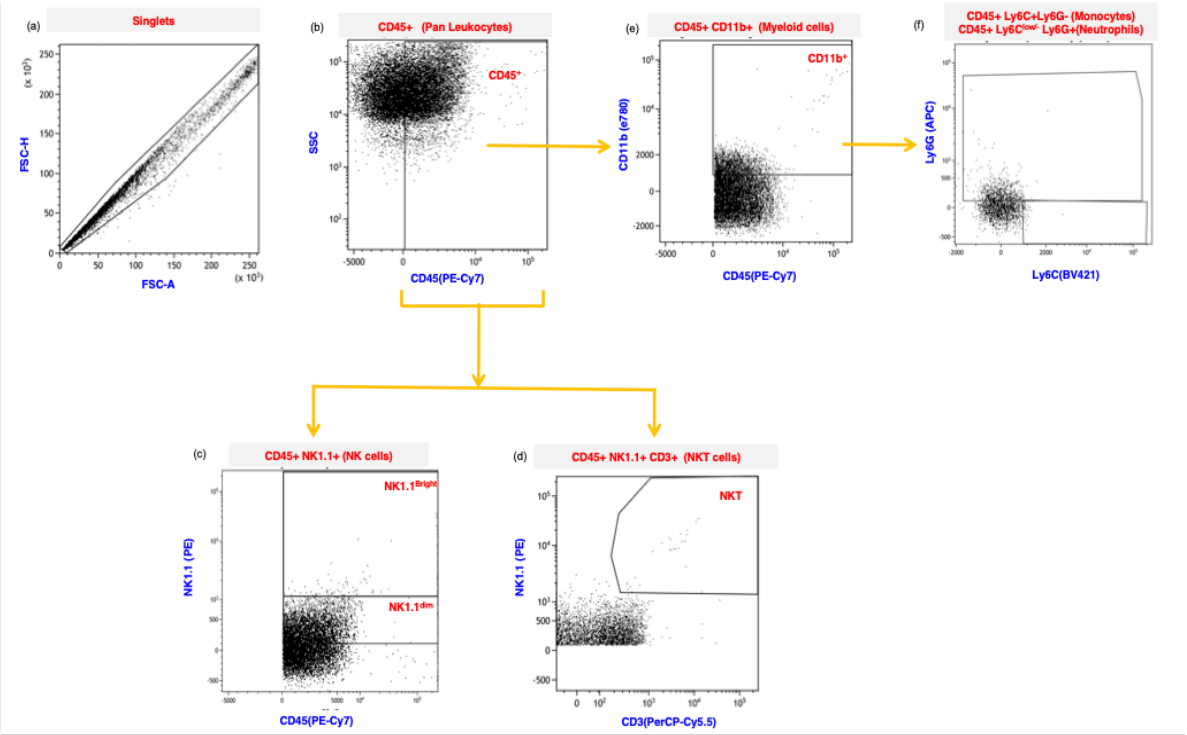


**Supplemental Figure 8 I** Gating strategy to identify NK cell subsets and Myeloid cell subsets in AH from cKO mice. Representative images show gating strategy used to determine the immune cell populations in cKO mice and control mice. (a, b) The marked region FSC-A versus FSC-H shows the singlet cells in the population and CD45 (APC-H7) versus Side Scatter (SSC) represents the leukocyte population. (c) This panel shows NK1.1 (PE) versus CD45(PE-Cy7), where double positive cells represent Natural Killer (NK) cells present among the CD45+ cells. The CD56 region is further subdivided into NK1.1$^{Bright}$ and NK1.1$^{dim}$ based on the expression of NK1.1 on cells. (d) The upper right panel represents Natural Killer T (NKT) cells that are positively stained for both CD3 (PerCP-Cy5.5) and NK1.1 (PE). (e) Top right region shows cells double positive for CD11b (e780) and CD45 (PE-Cy7) and represents the myeloid population within CD45+ cells.(f) The top region shows cells double positive for Ly6G (APC) and Ly6C(BV421), representing monocytes, while the bottom region shows cells positive for Ly6C (BV421) representing neutrophils. The percentages of immune cells were computed using number of positive cells for each marker to total number of cells acquired.

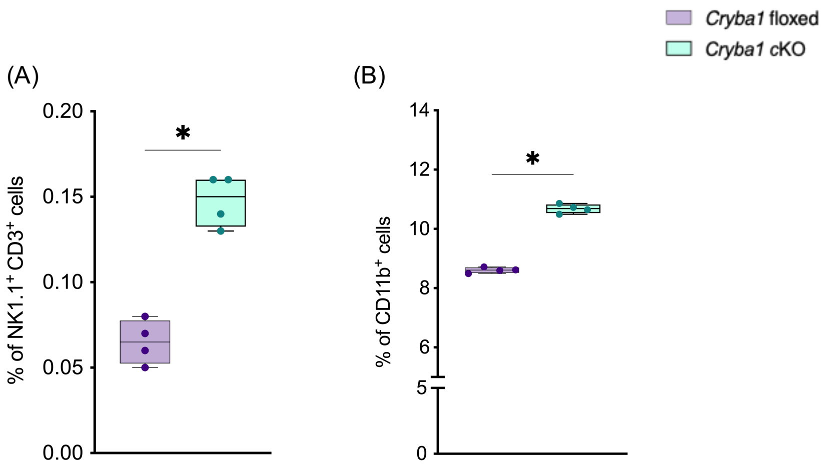


**Supplemental Figure 9 I** NKT cells and Myeloid cells in AH from cKO mice. (A,B) Box whisker plots show higher percentage of CD45+NK1.1+CD3+ (Natural Killer T – NKT cells) and CD45$^{+}$CD11b$^{+}$ (Myeloid cells) within total cells acquired from pooled aqueous humor of *Cryba1* floxed mice (n=4) and *Cryba1* cKO mice (n=4) at 15 months. Graphs show Min to Max (all points); *P < 0.05, Mann–Whitney test.

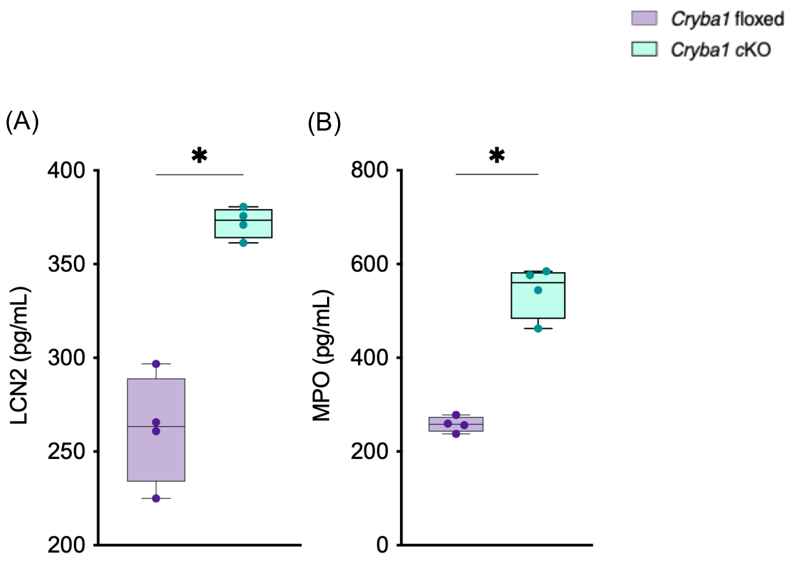


**Supplemental Figure 10 I** Increased levels of Neutrophil effector molecules in *Cryba1* cKO mice. ELISA showed increased LCN2 (A), and MPO (B) levels in *Cryba*1 cKO mice (n=4) at 15 months relative to *Cryba*1 floxed mice(n=4). Graphs show Min to Max (all points); *P < 0.05, Mann–Whitney test.

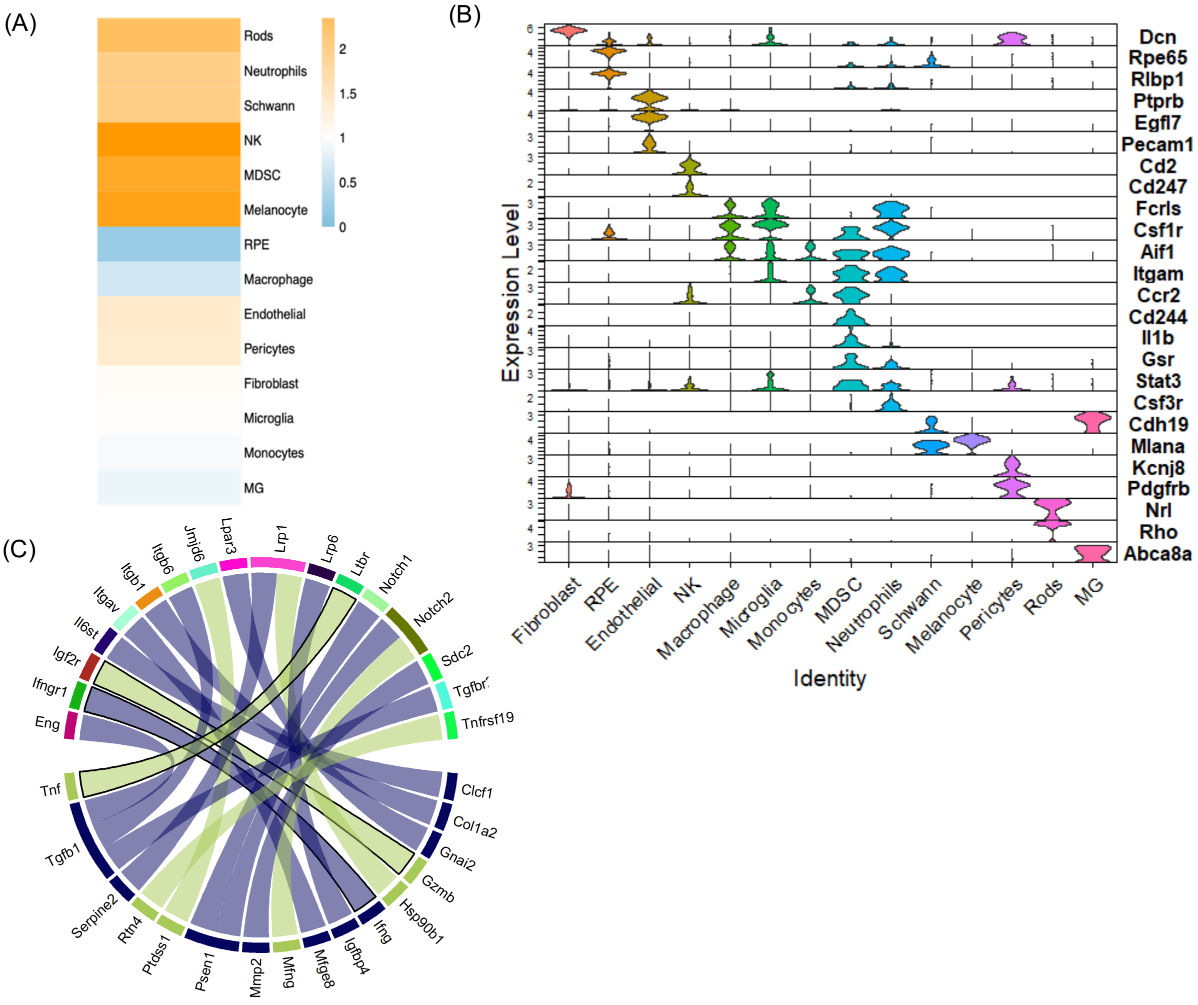


**Supplemental Figure 11 I** scRNA sequencing shows interaction patterns between NK-RPE cells in mouse SRS at 15 months. (A) Graph shows ratio of cell populations (cell pct) identified in *Cryba1* cKO and control mouse SRS with NK cells, neutrophils and MDSCs higher in *Cryba1* cKO mice. (B) Shows stacked vin plot of top marker genes identified in each cell population in mice SRS. (C) Chord plot of interaction partners curated from Cellinker with FC>1.3 and FC<0.8 in scores (*Cryba1* cKO versus control). The curved lines show relationships between the ligands identified for NK cells to receptors identified for RPE cells while colors of lines indicate known interaction partners (violet) curated and unique interaction partners (green) that not compiled by Cellinker. L-R interactions highlighted with black outline are those known in mediating cell death pathways

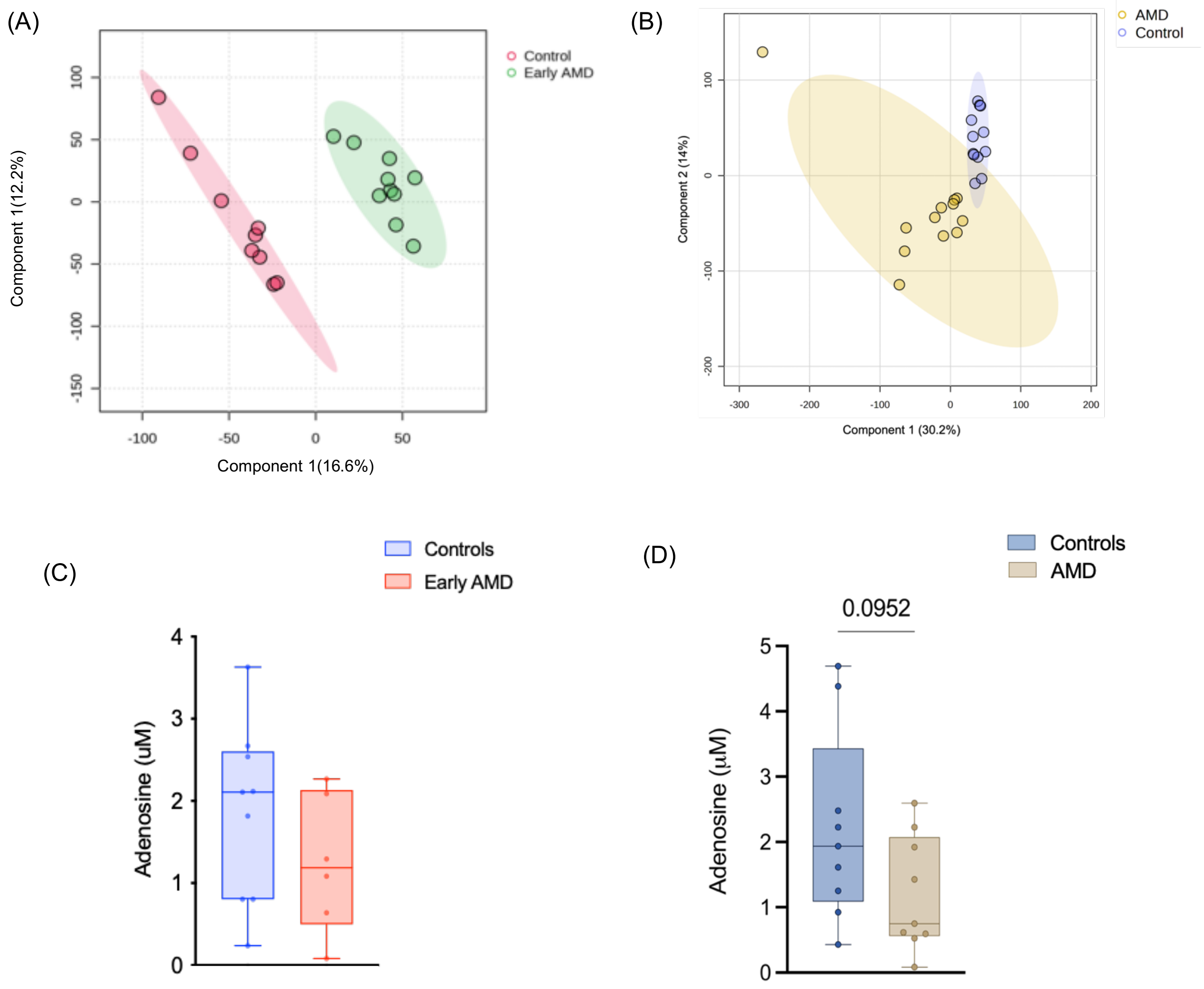


**Supplemental Figure 12 I** Metabolite analysis and adenosine assay of aqueous humor from human subjects. (A,B) Partial Least Squares Discriminant Analysis (PLS-DA) 2D plot of human metabolites in AH (patients) and retinal lysates (donor eye) of control and AMD. Components 1 and 2 show variability between groups indicating metabolic patterns are consistent within their respective groups. Shaded circles represent 95% confidence interval and the color circle indicate individual data points. (C) Graph shows adenosine assay of control and early AMD aqueous humor show decrease in adenosine concentration (μM) in early AMD (n=6) compared to cataract control (n=9). (D) Graph shows lower adenosine levels(μM) in retinal lysates in AMD donor retina compared to healthy retina (n=9) in both groups. Graphs show Min to Max (all points) obtained using Unpaired t test with Welch's correction.

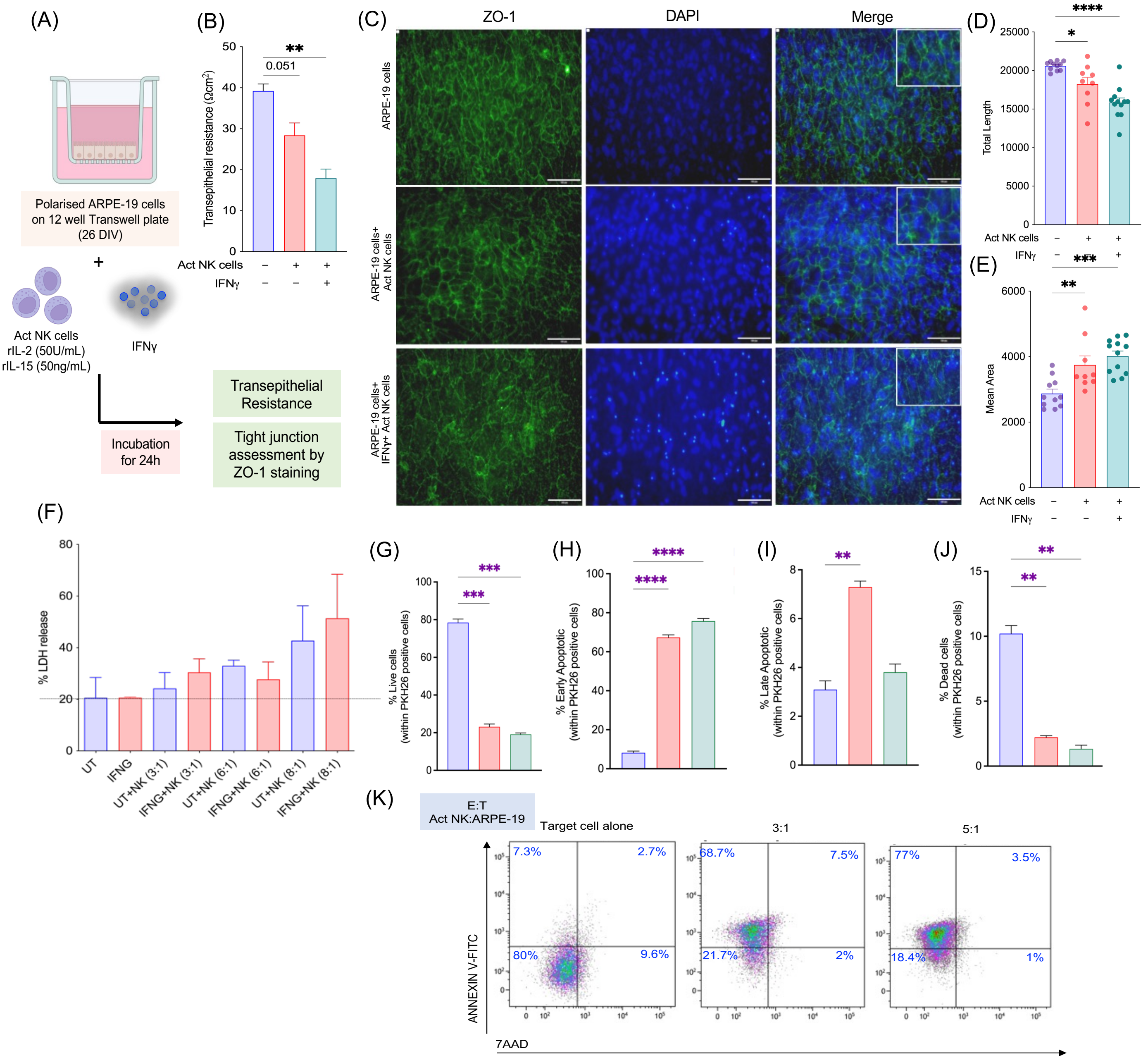


**Supplemental Figure 13 I** NK cell interaction impairs RPE barrier function and triggers death in ARPE-19 cells. (A) Methodology schema shows addition of activated NK cells and IFNγ to polarized ARPE-19 cells cultured in 12 well transwell plates. (B) TEER (Ωcm2) measurements of 25 day polarized ARPE-19 cells (+/-IFNγ) show significant reduction in presence of activated NK cells. (C) Immunofluorescence staining for Zonal occludin (ZO-1) in polarized ARPE-19 cells (control), ARPE-19 cells with NK cells, and ARPE-19 cells with NK cells and IFNγ (50ng/mL) at 20X magnification; ZO-1-Alexa fluor 488- Green, DAPI – nuclear stain(blue) in Olympus CKX53 microscope. Representative images are shown from cultures on addition of NK cells from 3 donors. Image J quantification shows significant reduction in (D) total length of network identified with increase in mean area (E) of the segments formed. (F) Control and IFNγ treated ARPE-19 cells co-cultured with activated NK cells subject to cytolysis at different ratios of effector (E) and target (T) cells ranging from 3:1, 6:1, 9:1 showed increased LDH activity irrespective of IFNγ addition. (G-K) PKH26 labelled control ARPE-19 cells (Target-T) and activated NK cells (Effector-E) were cultured at different E/T ratios(3:1 and 5:1) .Percentage of live cells, early apoptotic cells, late apoptotic cells and dead cells are shown as bar graphs, Reduced percentage of live and dead cells and increased percentage of early and late apoptotic cells are observed on co-culturing ARPE-19 cells and NK cells. PKH26 positive cells double negative for Annexin V-FITC and 7-AAD stain were marked as live cells, those positive for Annexin V-FITC were marked as early apoptotic cells, double positives for Annexin V-FITC and 7-AAD were marked as late apoptotic cells and 7-AAD stained cells were marked as dead cells. Graphs show Min to Max (all points); *P< 0.05, **P<0.01, ***P<0.001, ****P<0.0001 obtained using Ordinary one way ANOVA followed by Tukey's multiple comparison test (B,D,E,G-J).

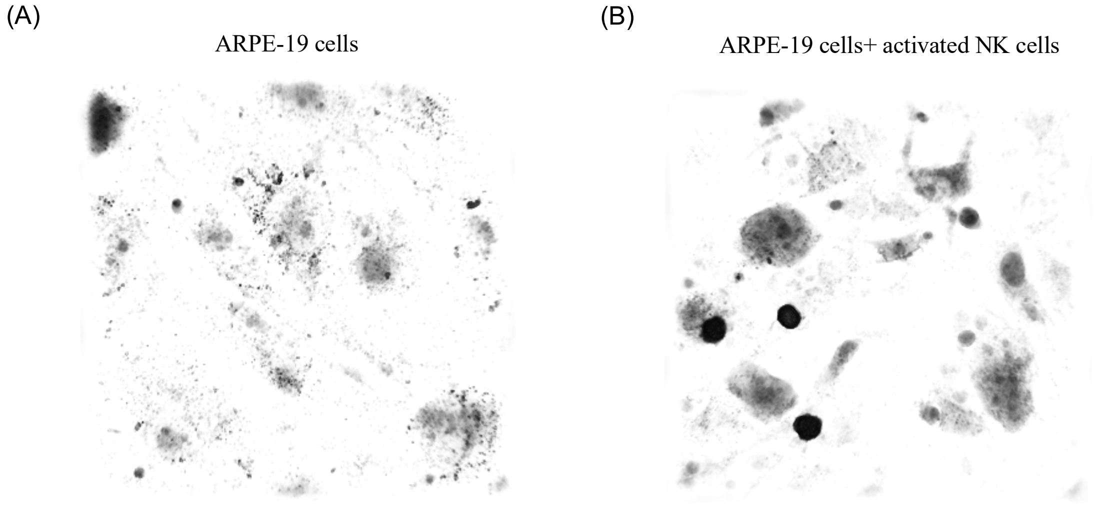


**Supplemental Figure 14 I** 3D Refractive index maps of ARPE-19 cells (A) and ARPE-19 cells co-incubated with activated NK cells (B) for 12 hours showing more dense cytoplasm in ARPE-19 cells on interaction with activated NK cells. Scale bar= 20 μm. Time lapse videos for all acquisitions were done at a rate of 5min/frame and data exported at 10fps. Scale bar =10μm

.

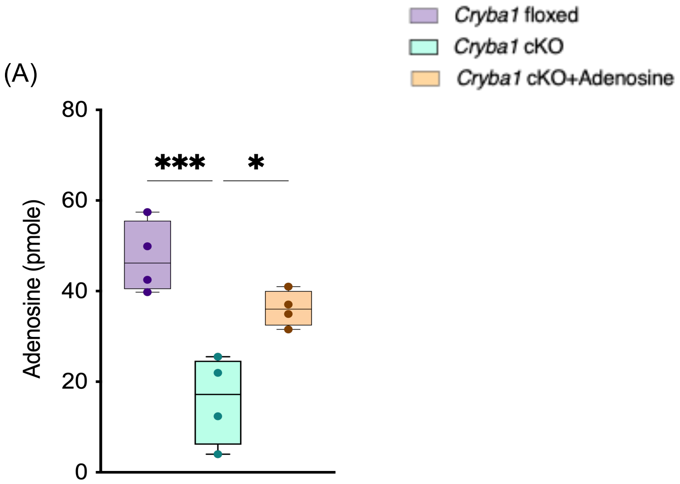


**Supplemental Figure 15 I** Intravitreal administration of adenosine improves adenosine levels in *Cryba1* cKO mice. (A) Adenosine quantity (pmole) assessed by ELISA was higher in *Cryba1* cKO mice treated with adenosine . Graphs show Min to Max (all points); *P < 0.05, ***P<0.001, Ordinary one way ANOVA followed by Tukey's multiple comparison test.

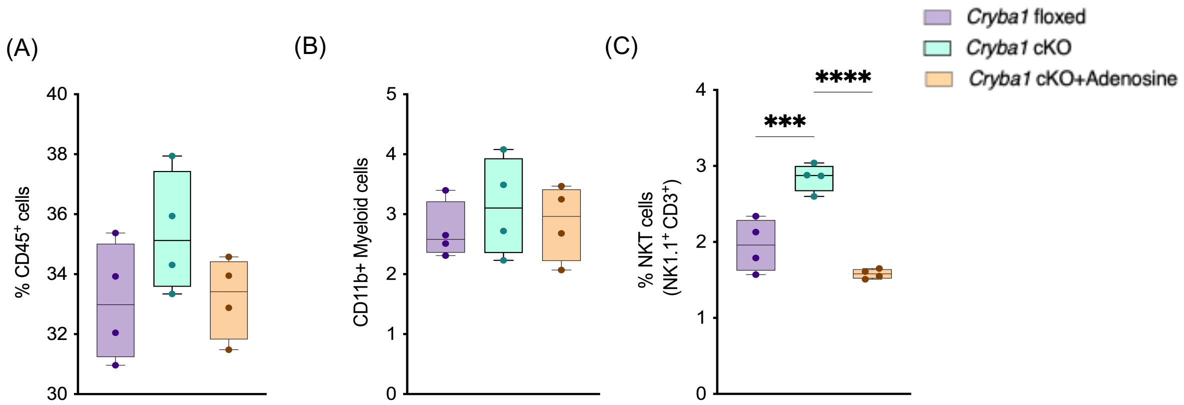


**Supplemental Figure 16 I** Altered proportions of leukocytes, NKT cells and myeloid cells in aqueous humor of cKO mice with adenosine injected intravitreally . (A-C) Box whisker plots show percentage of CD45+ (Leukocytes), $CD45^{+}CD11b^{+}$ (Myeloid cells) and CD45+NK1.1+CD3+ (Natural Killer T – NKT cells) within total cells acquired from pooled aqueous humor of *Cryba1* floxed mice, *Cryba1* cKO mice, and *Cryba1* cKO mice with intravitreal adenosine at 10 months (n=4 in each category). Graphs show Min to Max (all points); ***P<0.001, ****P < 0.0001, Ordinary one way ANOVA followed by Tukey's multiple comparison test.

### Supplemental Tables

**Supplemental Table 1 I** Aqueous humor soluble factor profiling in early AMD subjects. Table shows absolute levels of 48 soluble factors analyzed in aqueous humor of cataract (n=15) and early AMD (n=12) subjects. Values are shown as Average±SEM; *P < 0.05, **P < 0.01,***P < 0.001,****P < 0.0001, Mann–Whitney test and fold change between early AMD and cataract controls.

**Supplemental Table 2 I** L-R interactions in mouse curated based on Cellinker tool. Table shows known (A) and unique L-R interactions (B), ligand type and interaction type in mice retina based on fold changes in interaction scores of cKO versus control mice at 15months.

**Supplemental Table 3 I** The table lists pathways associated with L-R interactions and/or ligands from KEGG and Reactome pathway databases.

**Supplemental Table 4 I** The table highlights the interactions of NK ligands and RPE receptors involved in pathways related to programmed cell death from KEGG and Reactome pathway databases.

**Supplemental Table 5 I** Table shows the list of metabolites, mz values, retention time, peak intensity, fold change and VIP scores in control and early AMD aqueous humor.

**Supplemental Table 6 I** Table shows the list of metabolites, mz values, retention time, peak intensity, fold change and VIP scores in control and AMD donor retinal lysate.

**Supplemental Table 7 I** Table shows metabolomics profiling in retinae of control and cKO mice (4 months)

**Supplemental Methods**

**Antibody and reagents**

Fluorescent conjugated antibodies for human aqueous humor and donor eye immunophenotyping

Antibodies to CD45-APC-R700 (2μL/test, A79390), CD7- BV421 (4μL/test, 562635), CD16-APC-AF750 (5μL/test,A66330),CD159A-PC7 (5μL/test, B10246), CD11b-APC (4μL/test, A87782), CD15-FITC (7μL/test, IM1423U), HLA-DR-BV605 (4μL/test, IM3636) was obtained from Beckman Coulter, USA.

Antibodies to CD56-VioBRB515 (2μL/test,130-113-309), KIR2D-APC (2.5μL/test,130-092-687), Siglec 8-PE-Vio770 (5μL/test, 130-105-524) was procured from Miltenyi Biotec, USA.

Antibodies to NKG2D-PerCP-Cy5.5 (2.5μL/test, 562364), NKp44-PE (5μL/test, 558563), NKG2C-BV510 (5μL/test, 748167), CD14-PerCP-Cy5.5 (3μL/test, 562692), CD66b-PE (5μL/test, 561650), CD45-APC-H7 (2.5μL/test, 560178), CD3-PE (10μL/test, 555340), CD19-PerCP (10μL/test, 3475444), CD14-FITC (2.5μL/test, 347493), CD56-PE-Cy7 (2.5μL/test, 557747), CD66b-AlexaFluor 647 (2.5μL/test, 561645) was obtained from BD Biosciences, USA.

CD161-BV605 (5μL/test, 339916), CD69-BV510 (5μL/test, 310936), CD33- BV510 (4μL/test, 366610) were procured from Biolegend,USA.

Fluorescent conjugated antibodies for mouse aqueous humor phenotyping:

The antibodies include CD45-PE-Cy7 (1μL/test, 103114), anti-Ly6G-APC(1μL/test, 560599), Ly6C-BV421 (1μL/test, 562727), CD11b-AF700 (1μL/test, 557960) procured from BD Biosciences, USA; NK1.1-PE(1µL/test, 12-5941-82) procured from ThermoFischer Scientific, USA; CD27-FITC (1µL/test, 124207) and CD3-PerCP-Cy 5.5 (1µL/test, 100217) procured from Biolegend,USA

Antibodies for western blot analysis: Primary antibodies raised in rabbit include Gasdermin E antibody, 1:1000, 84005, Cell Signaling Technology,USA), anti GAPDH, (1:5000, 2118S, CST,USA) and secondary antibody used is Anti-rabbit IgG, HRP linked antibody(1:5000, 7074P, Cell Signaling Technology,USA).Primary antibody used for immunofluorescence is ZO-1 Monoclonal antibody (1:200, 339100, Invitrogen, ThermoFischer Scientific) and secondary antibody used is Goat Anti-Mouse IgG H&L (Alexa Fluor® 488) (1: 1000, ab150113, Abcam,USA). For mouse experiments, primary antibody used was mouse Gasdermin E (40618, Cell Signaling Technology)

Recombinant proteins/reagents used in the study includes recombinant human IL-2 carrier free (500U/mL, 589108,Biolegend, USA), recombinant human IL-15 carrier free (50ng/mL, 570308, Biolegend, USA), recombinant human IFNγ protein(50ng/mL, 285-IF-100, R&D Systems, USA), and Adenosine bioreagent, cell culture mammalian (5µM, A4036, Sigma-Aldrich, Merck, USA). Adenosine assay kit (MET-5090, Cell Biolabs, USA) was used for human subjects and mouse adenosine assay (ab211094, Abcam, USA) for mouse AH samples.

Cell culture media: DMEM-F12/L-Glutamine and RPMI 1640, fetal bovine serum (FBS), Antibiotic-Antimycotic (100X) were from GIBCO Invitrogen (Carlsbad, CA, USA).

**Processing of retinal samples from donor eyes for immunophenotyping and paraffin block preparation**

The whole globes were frozen using liquid nitrogen and dissected into equal half to extract the vitreous humor without the disturbing the retinal layers. One half was processed for preparing single cell suspension of retinal layers including RPE, but excluding the choroid. Single cell suspension was prepared by dissociating the tissue with accutase by agitation at 37°C and mechanical trituration in 1xPBS followed by washing by centrifugation post filtration with 40 μm cell strainer followed by immunophenotyping. The other half of the globe was formalin fixed and paraffin embedded block preparation was used for immunostaining and H&E staining.

**Aqueous humor sample collection in human subjects**:

Aqueous humor (AH) was collected intra-operatively at the time of cataract surgery by performing anterior chamber paracentesis. Briefly, 50-100 μL of AH was collected by inserting a syringe with a 30 mm gauge needle through the peripheral cornea. The AH was then centrifuged at 400g for 5 minutes at 4°C to separate cells for immunophenotyping and cell-free supernatant for multiplex ELISA. The cell-free supernatants were stored at -80°C till further use.

**Soluble factors measurement in human aqueous humor and cadaver vitreous humor**

The factors in aqueous humor were measured using Cytometric Bead Array (BD™ CBA Human Soluble Protein Flex Set System, BD Biosciences, USA) for the following analytes: IL-1α,IL-1β, IL-2, IL-6, IL-9, IL-10, IL-12/IL23p40, IL-12p70, IL-13, IL-17A,

IL-21, TNFα, IFNα, IFNγ, MCP1/CCL2,RANTES/CCL5, Eotaxin/CCL11, IL-8/CXCL8, MIG/CXCL9, IP-10/CXCL10, ITAC/CXCL11, Fractalkine/CX3CL1, TGFβ1, bFGF, VEGF, sICAM1, sVCAM, sL-selectin, sP-selectin, sTNFRI, sTNFRII, sIL-1R1, Angiogenin, IgE and sFasL were measured using as per manufacturer's instruction. Briefly, 50uL sample was incubate with 50uL capture bead cocktail for 1h to which 50uL detection reagent cocktail was added and incubated for 2h at room temperature with 500rpm agitation. The samples were washed and resuspended in CBA wash buffer.

Similarly, LIF, IL-18, IFNβ, HGF, EPO, PDGFAA, PDGF-BB, MMP2, MMP9, TIMP1, MPO, NGAL, Granzyme-B, Perforin, and β2microglobulin were measured using LEGENDplex™ (Biolegend Inc, USA) according to manufacturer's instructions. 25uL sample was incubated with 25uL capture bead mix for 2h at room temperature with 500rpm agitation. To this mix, 50uL detection reagent was incubated for 1h and 50uL of streptavidin-PE was incubated for 30minutes at room temperature with 500rpm agitation. The samples were washed and resuspended in 1X Biolegend wash buffer.

**Aqueous humor collection from mouse eye**

Mouse AH was collected using Harvard Pump Microinjection System and pulled glass micropipettes as explained previously(66). Briefly, the micropipette tip was broken at an angle and then inserted through the cornea into the anterior segment of the eye. AH was isolated through the pipette yielding about 4–5μL of AH per one eye.

**NK cell enrichment from healthy subjects**:

Healthy human peripheral blood (~20mL) was collected in 10mL BD vacutainer tubes with K2 EDTA (357525, Becton Dickinson, India). NK cell enrichment was done from freshly collected whole blood using EasySep$^{TM}$ Direct NK cell isolation kit (19665, Stemcell$^{TM}$ Technologies, USA) using manufacturers' protocol. Briefly, NK cells were enriched from whole blood using magnetic beads and antibody cocktails by negative selection. RBC lysis using NaCl (hypotonic lysis method) was employed to remove RBC contamination. The purity of NK cells post enrichment was assessed by staining with CD56 and CD3 antibody using flow cytometry (purity data not shown).
